# Integrated signaling and transcriptome analysis reveals Src-family kinase individualities and novel pathways controlled by their constitutive activity

**DOI:** 10.1101/2022.09.30.510317

**Authors:** Nikolaos Koutras, Vasileios Morfos, Kyriakos Konnaris, Adamantia Kouvela, Athanasios-Nasir Shaukat, Constantinos Stathopoulos, Vassiliki Stamatopoulou, Konstantina Nika

**Affiliations:** Department of Biochemistry, School of Medicine, University of Patras, Patras, Greece

**Keywords:** Src-family kinases, B-cell receptor signaling, ER-phagy

## Abstract

The Src family kinases (SFKs) Lck and Lyn are crucial for lymphocyte development and function. Albeit tissue-restricted expression patterns, the two kinases share common functions, the most pronounced one, being the phosphorylation of ITAM motifs in the cytoplasmic tails of antigenic receptors. Lck is predominantly expressed in T-lymphocytes; however, it can be ectopically found in B-1 cell subsets and numerous pathologies including acute and chronic B-cell leukemias. The exact impact of Lck on the B-cell signaling apparatus remains enigmatic and is followed by the long-lasting open question of mechanisms granting selectivity amongst SFK members. In this work we sought to investigate the mechanistic basis of ectopic Lck function in B-cells and compare it to events elicited by the predominant B-cell SFK, Lyn. Our results reveal substrate promiscuity displayed by the two SFKs, which however, is buffered by their differential susceptibility towards regulatory mechanisms, revealing a so far unappreciated aspect of SFK member-specific fine-tuning. Furthermore, we show that Lck- and Lyn-generated signals suffice to induce transcriptome alterations, reminiscent of B-cell activation, in the absence of receptor/co-receptor engagement. Finally, our analyses revealed a yet unrecognized role of SFKs in tipping the balance of cellular stress responses, by promoting the onset of ER-phagy, an as yet completely uncharacterized process in B-lymphocytes.

**Significance:** The Src-family-kinases Lck and Lyn are mandatory for lymphocyte function. However, several aspects of their regulation and critical pathways they control remain elusive. Using signaling and transcriptome analysis we show that the two kinases share substrate preferences; yet they display differential susceptibility towards regulatory mechanisms, revealing a so far unappreciated aspect of SFK member-specific fine-tuning. Furthermore, overexpression of both kinases suffices to induce receptor-ligation independent signaling responses. Finally, our analyses reveal a novel role of SFKs in tipping the balance of cellular stress responses, by promoting ER-phagy, in the expense of proteasomal degradation and the Unfolded Protein Response. These data advance our understanding of molecular individualities amongst SFK members, and identifies novel networks significant for lymphocyte activation and effector function.

## Introduction

Src-family kinases (SFKs) are ancient and evolutionarily conserved molecules. Their enzymatic activity is mandatory for several survival and activation-dependent biological processes in the context of many different tissues and cell types. Consequently, deregulated SFK activity has been described as an underlying cause for several neoplastic transformations, with the prototype of the family, c-Src, being the first ever described proto-oncogene (*1*).

SFK enzymatic efficiency is regulated by reversible phosphorylation events. Particularly relevant is a conserved tyrosine residue within their catalytic center, Y416 (Src numbering, used throughout this report). Trans-autophosphorylation of Y416 serves to stabilize the activation loop for optimal substrate binding and constitutes a conserved mechanism for SFK activation. Several protein tyrosine phosphatases (PTPs) have been attributed a role in keeping cellular levels of SFK activity at bay by dephosphorylating Y416 including CD45, SHP-1 and PTPN22 (*1*).

SFK members show ubiquitous and tissue-restricted expression, with Lck and Lyn being preferentially relevant for T- and B-lymphocyte function, respectively. Lck’s primary job is the phosphorylation of ITAM (Immunoreceptor Tyrosine -based Activation Motif) tyrosines located in the CD3 and ζ chains of the T-cell Receptor (TCR) complex and the consequent triggering of signalling responses after TCR engagement. Lyn performs a similar function on the B-cell Receptor (BCR) ITAMs (*1*).

Lck is found constitutively active in resting T cells, and the levels of the preactivated pool are necessary but also sufficient to transduce TCR-driven signalling responses without a need for further augmentation (*2*). Ectopic presence of Lck has been recorded in different types of malignant cells, however the functional relevance of this erroneous expression has almost exclusively been investigated in the context of CLL B-cells. Several studies have documented the concurrence between ectopically expressed Lck in CLL, and enhanced BCR-driven activation (*3*). Further to that, our previous work (*4*) designated Lck overexpression a marker of CLL B-cell subpopulations with heightened constitutive signaling profiles. Although these studies indicate the involvement of Lck in triggering autonomous BCR signals, several unmapped factors that can be present within neoplastic cells, prohibit the direct affirmation of a cause-effect relationship.

With the present work we sought to determine whether ectopic Lck overexpression could indeed ignite ligand-independent signaling reactions in B-cells, and if those would potentially differ from the conventional pathways triggered by the endogenous B-cell SFK, Lyn. Our data showed that alike Lyn, Lck can initiate autonomous B-cell signaling and alter the gene expression landscape of resting B-cells, albeit with a seemingly compromised efficacy. We further found that enzymatic promiscuity of SFKs can be buffered by their differential susceptibility to regulatory control mechanisms designed for keeping global SFK activity levels under strict control, thus preventing the initiation of unscheduled signaling. Finally, transcriptome analysis uncovered a yet unrecognized role of SFK signaling in cellular stress responses by favoring the onset of ER-phagy in the expense of proteasome-mediated degradation and the unfolded protein response (UPR).

## Results

### Ligation-independent BCR signaling is triggered by Lck overexpression

To enable studies that would provide evidence for a direct impact of Lck on BCR signaling pathways, we developed a stable B-cell line model. We used the Burkit lymphoma cell line, BJAB and introduced Lck via lentiviral gene transfer. To avoid counterselection and/or signaling pathway counterbalance events, often occurring as a result of the stable expression of signaling mediators in cell lines, we utilized a Doxycycline (Dox)-inducible system for Lck expression. Lck was invariably untraceable in non-Dox treated cultures (designated -Dox henceforth). Supplemental Text and Fig. S1 describe selection criteria and characterization of the B-cell model system. Similar to its physiological T-cell environment, Lck was also constitutively active within BJAB cells. Importantly, the introduction of Lck in BJAB significantly increased the baseline levels of SFK activity (Fig. 1A upper panel, and S2A) as determined by staining with a-pY416, an antibody specifically recognizing the phosphorylated form of the activation loop tyrosine of SFKs and thus widely used as an indicator of pan-SFK activity (*2, 5*). The first step of B-cell activation from the BCR, is the phosphorylation of the Igα and Igβ ITAM tyrosines by Lyn (*6*). Lck performs exactly the same job on the TCR ITAMs (*3*). Staining with an Ab recognizing the phosphorylated Igα ITAM Y182 (designated pCD79a henceforth) revealed that Lck expression in BJAB resulted in BCR ITAM phosphorylation without a requirement for receptor ligation. pCD79a was abrogated after treatment with a specific Lck inhibitor, Lcki, or by expression of LckΔSH4, a mutant lacking the moiety responsible for anchoring at the inner leaflet of the plasma membrane (*3*). Therefore, BCR ITAM phosphorylation required Lck’s intact enzymatic activity and its juxtaposition to the receptor (Fig. 1A).

**Figure 1.**
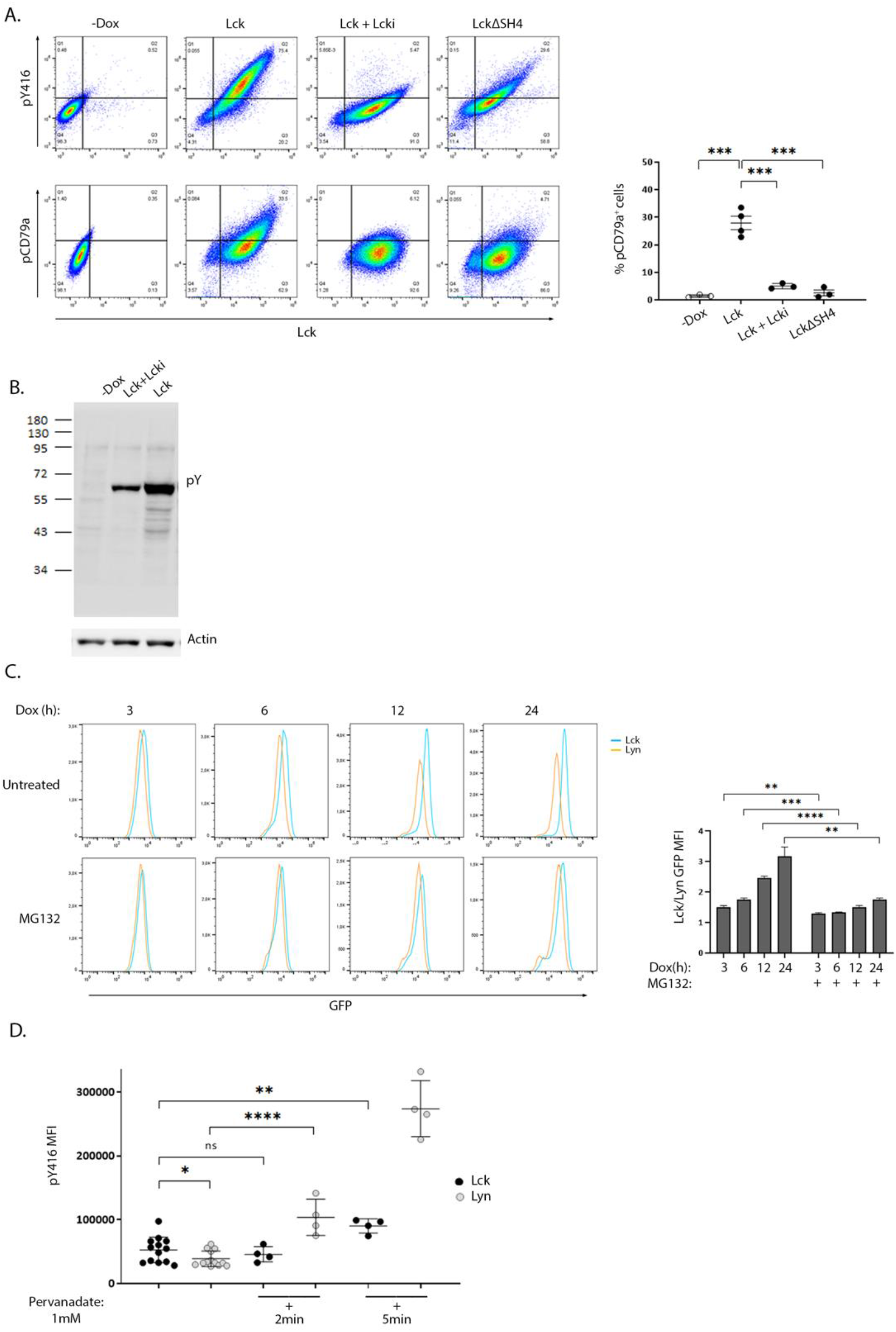
Differential susceptibility to regulatory control mechanisms, sets the threshold for SFK activity and ligation-independent BCR signaling. **A. High expression of ectopic Lck in B-cells drives constitutive BCR ITAM phosphorylation**. BJAB cells transduced with LckWT or the deletion mutant ΔSH4 were cultured in the absence or presence of Dox for 48h. LckWT-expressing cells were further treated with 10 μM of the specific Lck inhibitor Lcki for 10 min. Samples were stained for Lck, its’ active form (pY416) or pCD79a and are presented as 2D FACS plots of Lck^+^ gated cells, to better visualize the dose-response relationship between Lck expression and its active form (upper panels) or ignition of constitutive BCR ITAM phosphorylation (lower panels). Quadrant gating was set on the corresponding negative populations (-Dox samples). The right-hand panel graph shows the frequency of cells in the upper right quadrant (double-positive for Lck and pCD79a) (n≥3, Unpaired Student t test; mean +/-standard deviation [SD]; ***P < 0.001). **B. Ectopic Lck in B cells ignites signaling pathways downstream the BCR** Anti-phosphotyrosine (pY) clone 4G10 western blot of total cell lysates from LckWT-expressing cells untreated or incubated with 10 μM of Lcki for 10 min and the -Dox treated sample. Actin blot (lower panel) verifies equal sample loading. **C. Lyn shows significantly higher predisposition to proteasomal degradation than Lck** Cells expressing GFP-tagged Lck or Lyn were either left untreated or incubated with 1 μM of the proteasome inhibitor MG132. Samples were removed from the culture at the indicated time points after Dox addition and were analysed by FACS to monitor SFK protein production via GFP fluorescence intensity. Graph shows the ratios of Lck/Lyn GFP Mean Fluorescence Intensities (MFIs) in the absence or presence of MG132 at each indicated time point (n=3, Unpaired Student t test; mean +/- SD; **P < 0.01, ***P < 0.001, ****P < 0.0001). **D. The phosphorylation status of the activation loop tyrosine of Lyn is under stricter control by PTPs** Cells expressing GFP-tagged Lck or Lyn were either left untreated or incubated with 1mM Pervanadate for the indicated times. Graph shows pY416 MFI values within cell populations gated for comparable SFK expression levels (equal GFP fluorescence) (n=13 for untreated and n=4 for pervanadate treated cells, Unpaired Student t test; mean +/- SD; *P < 0.05, **P < 0.01, ****P < 0.0001; ns: not significant).

2D FACS plots showed a dose-response relationship between Lck expression/activity and pCD79a induction and exposed a prerequisite for surpassing an expression threshold for receptor-independent ITAM phosphorylation to occur. Further to pCD79a, introduction of Lck to BJAB cells increased basal levels of global tyrosine phosphorylation, again in the absence of receptor or co-receptor ligation (Fig. 1B) implying that Lck’s effect on the BCR ITAMs is “productive” enough to propagate to downstream pathways. Importantly, these phosphorylation events were unrelated to Lyn or other SFKs, since we could not detect any shifts on their activation status, unpredictably resulting from the ectopic presence of Lck (Fig. S2B). Unlike BJAB cells, Lck overexpression in HEK293T failed to alter the global phosphotyrosine landscape, whereas Src and to a lesser extend Lyn and Fyn did (Fig. S2C), indicating a preference of Lck for antigenic receptor signaling.

### Lck and Lyn differ in the efficiency for signal ignition and in their susceptibility to regulatory mechanisms in B-cells

In light of these observations, we asked whether ignition of autonomous signaling results from failure of B-cell regulatory mechanisms to restrain an “alien” kinase, or if this is a general outcome of increased SFK activity levels, overruling the processing capacity of inhibitory pathways. To answer this conundrum, we deduced that a comparison between ectopic Lck and “resident B-cell SFK” Lyn, would prove informative. This type of analyses required a means of readily comparing expression levels between the two SFKs. We thus introduced a GFP-tag on their C-termini. When expressed in BJAB cells, the GFP-tagged proteins behaved identically to their non-GFP versions, with regard to pY416 levels and their ability to induce constitutive pCD79a (Fig. S3A), confirming that the presence of GFP did not interfere with their biological function and thus the tagged constructs could be reliably used for the continuation of our studies. To obtain cell populations with homogeneous levels of SFK expression/activity, cell lines were sorted for GFP^+^ cells (Fig. S3B) and all subsequent analyses were performed in the sorted GFP^+^ populations.

During the course of these experiments, we faced an insuperable difficulty to obtain Lyn expression levels equally high to those of Lck (Fig. S3B and S3C). This appeared to be a problem with Lyn protein stability within the lymphocyte environment since transient transfection of the two SFK constructs in HEK 293T cells resulted in comparable expression (Fig. S3D). The longer splice variant of Lyn, LynA, is particularly susceptible to ubiquitin-mediated breakdown, thus we used the degradation-resistant isoform (LynB), lacking the region which flags the kinase for Cbl-mediated polyubiquitination (*6*). However, even LynB could not reach expression levels comparable to those of Lck, a trend that was reversed after treatment with the proteasome inhibitor MG132 (Fig. 1C). Thus, the proteasomal degradation machinery constrains the abundance of Lyn, but not Lck, within B-cells.

Nonetheless, the presence of the GFP tag allowed gating and direct comparison of cell populations with equal levels of protein expression (Fig. S3E). Similar to Lck, Lyn was also constitutively active in B cells, as confirmed by pY416 staining, and its expression augmented SFK activity levels (Fig. S3C). However, the BJAB-Lck line not only contained higher amounts of global SFK activity, due to the surplus in Lck protein expression (Fig. S3C), but also higher pY416 when compared with Lyn on an equivalent protein expression basis (cells gated for equal GFP) (Fig. 1D). 2min treatment with the Tyrosine Phosphatase (PTP) inhibitor Pervanadate sufficed to drastically increase Lyn-pY416 whereas Lck-pY416 remained unaffected. Appreciable Lck-pY416 increases required much longer incubation times, at least 5-10 min of pervanadate treatment (Fig. 1D). This differential sensitivity of the two SFKs towards pervanadate strongly suggests that Lyn is subjected to stricter PTP control on the phosphorylation dynamics of its activation loop tyrosine.

Following antigen recognition and ITAM phosphorylation, propagation of BCR signaling requires the recruitment and activation of Syk and subsequent phosphorylation of its substrates. BCR ligation also activates the PI3K/Akt pathway, which is essential for B-cell function (*7*). Dox-induced expression of both Lck and Lyn drove the phosphorylation of signal transducers pCD79a, Syk and Akt, increased the global phosphotyrosine content in unstimulated BJAB cells (Fig. 2A and S4B) and provided a relative enhancement in BCR-triggered responses after a-IgM stimulation (Fig. S4). Consistent with the higher content of SFK activity, the phosphorylation fingerprint of downstream signaling was significantly more intense in the BJAB-Lck line. However, analysis of cells with equivalent SFK protein expression (equal GFP gates), revealed the superior efficiency of Lyn in BCR signaling, since Lck had to be expressed in much higher amounts in order to elicit responses of comparable magnitude (Fig. 2B).

**Figure 2.**
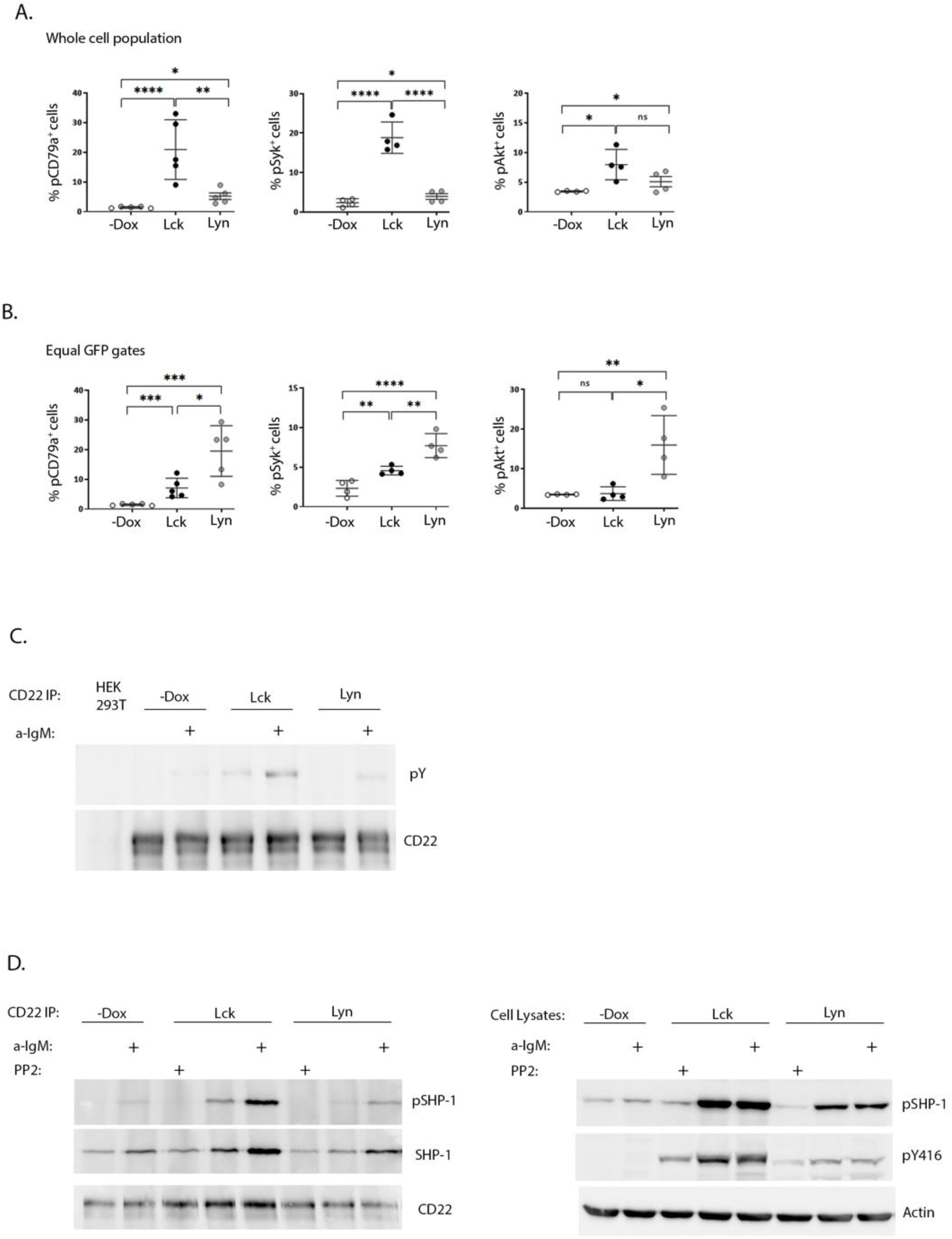
SFK overexpression suffices to drive ligation-independent positive and negative signaling pathways. **A. Lck-expressing cells have more robust signaling responses on a whole cell population basis** Resting Lck- or Lyn-expressing cells were stained for pCD79a, pSyk (Y525/526) and pAKT (Ser473) and analysed by FACS. Graphs display the frequency of cells double-positive for GFP and pCD79a, pSyk and pAKT respectively (Gating as in Figure 1A, % of cells in the upper right quadrants) (n≥4, Unpaired Student t test; mean +/- standard deviation [SD]; *P < 0.05, **P < 0.01, ***P < 0.001, ****P < 0.0001; ns: not significant). **B. Lyn is more efficient than Lck in igniting phosphporylation of BCR signaling mediators on an equal protein expression level** Same as in A, but for equivalent levels of SFK expression (Fig. S3E for description of equal GFP gating strategy). **C. Enhanced CD22 tyrosine phosphorylation in response to SFK overexpression** CD22 was immunoprecipitated from the Lck- and Lyn-BJAB cell lines and their -Dox counterparts in the absence or presence of stimulation with 10 μg/mL a-IgM for 10min. IPs were analysed by western blotting with a-pY 4G10 (upper panel) and a-CD22 (lower panel) antibodies. HEK293T cells (devoid of CD22 expression) were used as a negative control for the IPs. **D. SFK overexpression ignites the CD22/SHP-1 inhibitory pathway and this depends on their intact kinase activity** <u>Left-hand panels</u>. CD22 IPs as in C, but with additional samples treated for 10 min with 100 μM of the pan-SFK inhibitor PP2, as indicated. IPs were analysed by western blotting with a-pSHP-1 Y564 (upper panel), total a-SHP-1 (middle panel) and a-CD22 (lower panel) antibodies. <u>Right-hand panels</u>. Total cell lysates from the same experiment, analysed by western blotting with a-pSHP-1 Y564 (upper panel) and a-pY416 (middle panel) to verify the PP2-induced reduction in SFK activity. Actin blot (lower panel) attests equal sample loading. Shown are representative blots of two independent experiments.

Collectively these data show that the BCR signaling machinery is more responsive to the action of Lyn, at the same time imposing stricter regulation on its expression and activity levels. On the other hand, the case of Lck shows that despite reduced efficiency for the B-cell signaling apparatus, when global SFK activity levels surpass a certain threshold, autonomous activation can occur. Therefore, SFK protein individualities and the total amount of SFK activity within a cell, are equally contributing factors in fine-tuning signaling thresholds for antigenic receptors.

### Constitutive triggering of the CD22/SHP-1 inhibitory axis by Lck and Lyn overexpression

Further to their necessity for ITAM phosphorylation Lyn, and to a lesser extent Lck, are capable of triggering parallel inhibitory pathways. CD22 is an extensively characterized inhibitory receptor in B-cells. CD22 ITIM phosphorylation by Lyn, creates docking sites for the recruitment of the PTP SHP-1 (*6*). Furthermore, Lck and Lyn can phosphorylate Y564 of SHP-1 which results in enhanced phosphatase activity (*6, 8*). Dox-induced expression of both SFKs was followed by enhanced constitutive and a-IgM induced phosphorylation of CD22 and SHP-1Y564 (Fig. 2C and 2D). Accordingly, stronger co-immunoprecipitation of CD22 with SHP-1 (and pSHP-1) was detected in SFK-expressing BJAB cells compared to the -Dox counterpart. This association, as well as pSHP-1 phosphorylation receded to baseline levels upon treatment with the pan-SFK inhibitor PP2 (Fig. 2D), demonstrating the interdependence of intact SFK kinase activity and upregulation of the CD22/SHP-1 inhibitory axis. Therefore, in addition to igniting signaling downstream the BCR, ectopic Lck can trigger ITIM-induced responses from CD22.

### SFK overexpression modifies the Transcriptome profile of B-cells

Further to their influence on proximal BCR signaling, we investigated the net-result of SFK-triggered pathways on the gene expression program. RNA extracted from BJAB lines overexpressing Lyn or Lck, 48h after Dox addition, and the non-Dox-treated counterparts were used to produce corresponding cDNA libraries for NGS and Bioinformatics analysis. Differentially expressed genes (DEG) were selected using two criteria: log fold change >0.5 or <-0.5 (for upregulated and downregulated DEGs respectively) relative to the -Dox sample, and p<0.05. We identified 1264 and 956 differentially expressed genes in the BJAB-Lyn and BJAB-Lck cell lines, respectively (GEO Accession number: GSE212759). Lyn overexpression coincided with upregulation of 759 genes and Lck with 517. 321 upregulated genes were common for the two SFKs. Likewise, the expression of 505 and 439 genes was repressed as a consequence of Lyn and Lck overexpression respectively, 219 of which were commonly downregulated by both SFKs (Fig. 3A and 3B).

**Figure 3.**
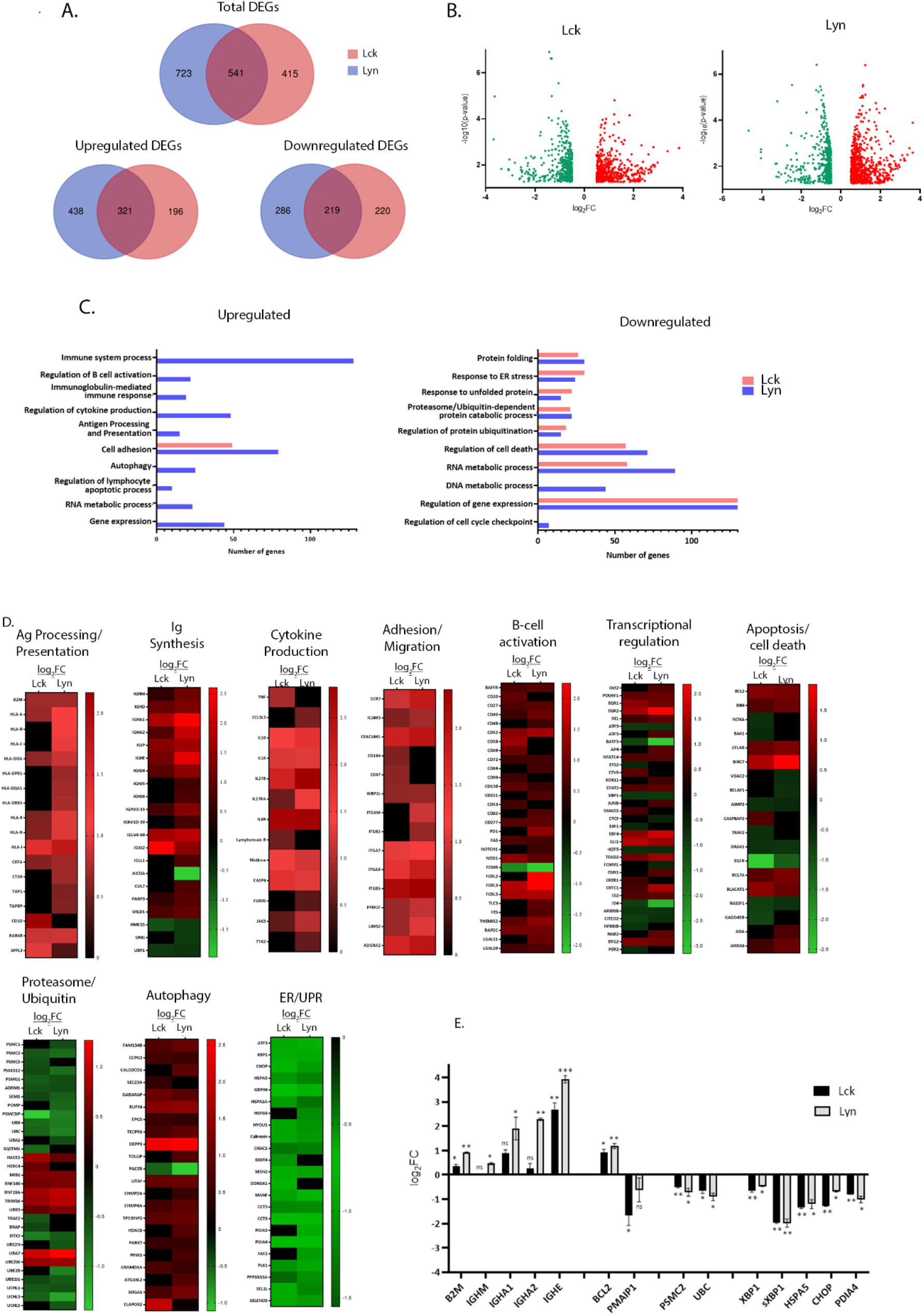
SFK overexpression suffices to change the gene expression landscape of unstimulated BJAB cells. **A. Lck and Lyn alter the expression profile of common and unique genes** Venn diagram representation of the RNAseq data from Lck- (pink) and Lyn- (blue) expressing cells. Top panel, number of total differentially expressed genes (DEGs). Lower panels, number of upregulated (left) and downregulated (right) DEGs. The numbers in the common intersecting areas designate overlapping DEGs. Thresholds for DEG selection: log_2_ fold change (log_2_FC) >0.5 and <-0.5 for upregulated and downregulated genes, respectively and adjusted p-value <0.05 (calculated using limma). **B. Distribution of upregulated versus downregulated DEGs in SFK overexpressing cells** Volcano plots of DEGs from Lck- (left) and Lyn-expressing cells (right). The graphs present the relationship between log_2_FC, and statistical significance (-log_10_(p-values)), using a scatter plot view. **C, and D. Lck and Lyn modulate DEG signatures assigned to the regulation of several biological processes C**. Graphical representation of Gene Ontology (GO) enrichment analysis of upregulated (left) and downregulated (right) DEGs in Lck-BJAB (pink) and Lyn-BJAB (blue) cells. The bar graphs illustrate the most representative enriched (FDR (false discovery rate) < 0.05) biological function classes and the number of genes corresponding to each GO term. **D**. Heatmaps illustrating transcript abundance (log_2_FC) for DEG sets involved in the indicated biological processes. Downregulated genes are marked green and upregulate red. Black denotes genes that are not differentially expressed. (n = 2 for each cell line). **E. RT-qPCR based validation of representative RNA-seq derived DEGs**. The y-axis shows the log_2_FC of the gene expression between Lck-BJAB (black)/-Dox and Lyn-BJAB (grey)/-Dox (n=2, Multiple unpaired Student t test; mean +/- SD; *P < 0.05, **P < 0.01, ***P < 0.001; ns: not significant).

Gene ontology (GO) enrichment analysis assigned differentially expressed genes to several functional outcomes ranging from processes reminiscent of B-cells that have received BCR engagement signals (B-cell activation, Ig and cytokine synthesis, Ag presentation) to regulation of apoptosis, autophagy and the ER stress response (Fig 3C.). Unique genes upregulated or downregulated by each SFK could not be assigned to any specific biological function by GO enrichment analysis, so at this point we cannot deduce that there is discrimination on the signaling pathways originating from Lyn or Lck overexpression. Although GO analysis of Lck-induced genes did not reveal any significant enrichment in biological processes other than cell adhesion/migration, a more thorough inspection showed Lck’s involvement in the same biological processes as Lyn (Fig. 3D and 3E).

SFK overexpression was followed by induction of class I and II HLA genes, the transcriptional co-activator CIITA and accessory molecules (CTSH, TAP1, TAPBP, RAB4B and SPPL3) essential for the formation and presentation of pMHC complexes (Fig. 3D, corresponding heat map). Several genes encoding constant and variable Immunoglobulin regions were significantly induced (Fig. 3D, corresponding heat map). In addition to the two BCR isotypes, IgHM and IgHD, mRNAs encoding the constant regions of IgA and IgE were strongly upregulated. Counterintuitively, expression of Activation-induced cytidine deaminase (AICDA/AID), the key initiator of class switch recombination (CSR), was repressed whereas the E3 Ubiquitin ligase responsible for AID protein degradation, CUL7 (*9*) was upregulated. Concomitantly, DNA repair factors acting alongside AID were differentially expressed. PARP3 a negative CSR regulator (*10*) was upregulated, whereas UNG and HMCES, both of which are required for efficient CRS (*11, 12*), were downregulated. Notably the transcriptional activator UBP1 (LBP-1a) was repressed. Suppression of UBP1 has been shown to preferentially promote IgA (but not IgG1) in B-lymphocytes induced to undergo CSR (*13*), which interestingly coincides with increase of IgHA but not IgHG mRNA levels.

Consistent with B-cells’ recognized contribution to the cytokine milieu, several cytokine encoding genes were induced together with the cytokine-activator member of the Caspase-1 family, CASP-4 (Fig. 3D, corresponding heat map).

The expression level of genes encoding proteins involved in cell migration and adhesion were also strongly upregulated (Fig. 3D, corresponding heat map). Amongst those were members of the integrin family and their regulators (PPM1F, LIMS2/Pinch-2), adhesion molecules (CEACAM1, ICAM3) and the adhesion G-protein coupled receptor ADGRA2, as well as the chemoattractant receptor CCR7, upregulation of which is one of the earliest events following BCR engagement (*14*).

Numerous B-cell co-stimulatory receptors, activation markers and signaling mediators (*15*) were also induced (Fig. 3D, corresponding heat map). Amongst those were, BAFFR (TNFRSF13C), one of the major pro-survival receptors in B-cells, CD40, CD48, CD69, CD52, CD20, CD27, the ITIM containing CD72 and the BCR-induced exhaustion marker, PD1 (*16*) as well as members of the FCRL family. mRNA levels of the cell-death receptor FAS were also upregulated, a feature characteristic of activated B-cells (*17*).

Expression profiles of several B-cell activation-relevant transcription factors and transcriptional regulators (*18*) were altered as a result of SFK-triggered signals in BJAB cells (Fig. 3D, corresponding heat map). Induced genes common between the two SFKs included ID2, an E2A and Pax5 antagonist, the EBF family member EBF4, the AP-1 transcriptional coactivator CRTC1, the NF-kB subunit REL (in parallel to downregulation of the NF-kB inhibitor, NF-kBIB), STAT2, SOX11 as well as the early growth response genes EGR1 and EGR2. BTG2, a transcriptional coregulator important for B-cell development and differentiation was also upregulated. In parallel, we detected repression of pro-apoptotic mediator/tumor suppressors ID4 and ARID5B, downregulation of which correlates with enhanced acute lymphoblastic leukemia (ALL) cell proliferation (*19*). The Lyn-specific signature comprised induction of Oct-2, ATF5, the AP-1 member JUNB, AP4 and NAB2 (a BCR-induced transcriptional co-activator of IL-2 production), with concomitant repression of CTCF (a BCR-induced growth suppressor). Lck-expressing cells preferentially upregulated CREB1, and SMAD2, an activator of IgA class switching. There appeared to be opposing regulation of two TFs belonging to the ETS family, namely downregulation of ETS-2 (an IL-2 mRNA repressor) and upregulation of EBV6 (a PAX5 suppressor). mRNA levels for E4F1, an inhibitor of cell proliferation and PER2, a negative regulator of lymphoid immune functions, were uniquely downregulated in response to Lck expression in BJAB cells.

Alterations in the balance between pro-survival versus pro-apoptotic transcriptome landscape appeared more complex (Fig. 3D, corresponding heat map). SFK-triggered pathways led to the induction of inhibitors of apoptosis BIRC7 and CFLAR. Repressed genes included AIMP2 an enhancer of TNF-mediated cell death and the apoptosis-promoting molecules DRAK1 (a member of the death-associated protein kinases -DAPK) and the transcriptional repressor BCLAF1. Expression patterns of genes belonging to the BCL2 family revealed upregulation of the key anti-apoptotic regulator Bcl2 and pro-apoptotic protein Bim whereas in Lck-expressing cells mRNA levels of the pro-apoptotic mediators NOXA and BAK1 were downregulated. Lyn expression was accompanied by repression of the VDAC2 gene, encoding for a voltage-dependent anion channel responsible for maintaining BAK1 in an inactive state.

Expression of several E3 Ubiquitin ligases were induced upon SFK overexpression, whereas ubiquitin modifiers and conjugating enzymes as well as mRNAs encoding for ubiquitins B and C were downregulated. Similar pattern of repression was observed for genes encoding a number of proteasome subunits and proteasome assembly and maturation proteins (POMP, PSMG1) (Fig. 3D, corresponding heat map).

A number of positive regulators of autophagy were induced. Amongst them were TP53INP2, EPG5, RUFY4 and TOLLIP whereas mRNA levels of autophagy enhancer RUBCNL/PACER were downregulated (Fig. 3D, corresponding heat map). Notably, genes encoding the ER-phagy receptors RETREG1/FAM134B, CCPG1 and CALCOCO1 and facilitators GABARAP and SEC23 were also induced.

Rather unexpectedly a significant proportion of repressed genes were related to the endoplasmic reticulum (ER) stress and the Unfolded Protein Response (UPR) pathways (Fig. 3D, corresponding heat map). These included a wide range of ER chaperones and enzymes (PDI4, HSPA1, HSPA9), UPR target genes and downstream regulators (HSPA5, GRP94, CHOP, CHAC1, SESN2, HERPUD1 and others), as well as UPR-related transcription factors XBP1, ATF3.

Altered mRNA levels of key genes discovered by the transcriptome analysis were validated by qPCR (Fig. 3E). Importantly we verified that the recorded gene expression alterations were unrelated to the addition of Dox to the cell cultures (Fig. S5).

### A novel role of SFKs in ER homeostasis

The abundance of downregulated genes assigned to ER homeostasis responses and given the central role of the endoplasmic reticulum on the B-cell secretory pathway, prompted us to investigate whether SFK-induced signals would influence processes related to the excessive demand for protein synthesis and folding. BJAB-Lck and BJAB-Lyn cells were removed from the culture at regular time intervals ranging from 0-48h after Dox addition, stained with ER tracker and analysed by FACS, at the same time monitoring SFK expression by GFP fluorescence intensity (Fig. 4A). Increases in SFK expression levels correlated with ER expansion. Importantly, ER size receded to basal levels after 18h incubation with the Src kinase inhibitor PP2, affirming that the observed ER expansion was a consequence of SFK-induced signals. Aliquots of cells stained with ER tracker for each corresponding time point, were analysed by qPCR for the expression of UPR mediators (Fig. 4B). It has been documented that XBP1, sXBP1, HSPA5 and CHOP mRNA levels significantly increase by 6h of chemically induced ER stress and remain elevated after to 12h (*20*). However, at no point we could detect upregulation of any of the four genes, despite the recorded ER expansion. These data indicate augmented protein load reaching the ER, forcing it to increase its volume in order to meet demands, but surprisingly this process seems to be accompanied by a parallel blocking of the UPR.

**Figure 4.**
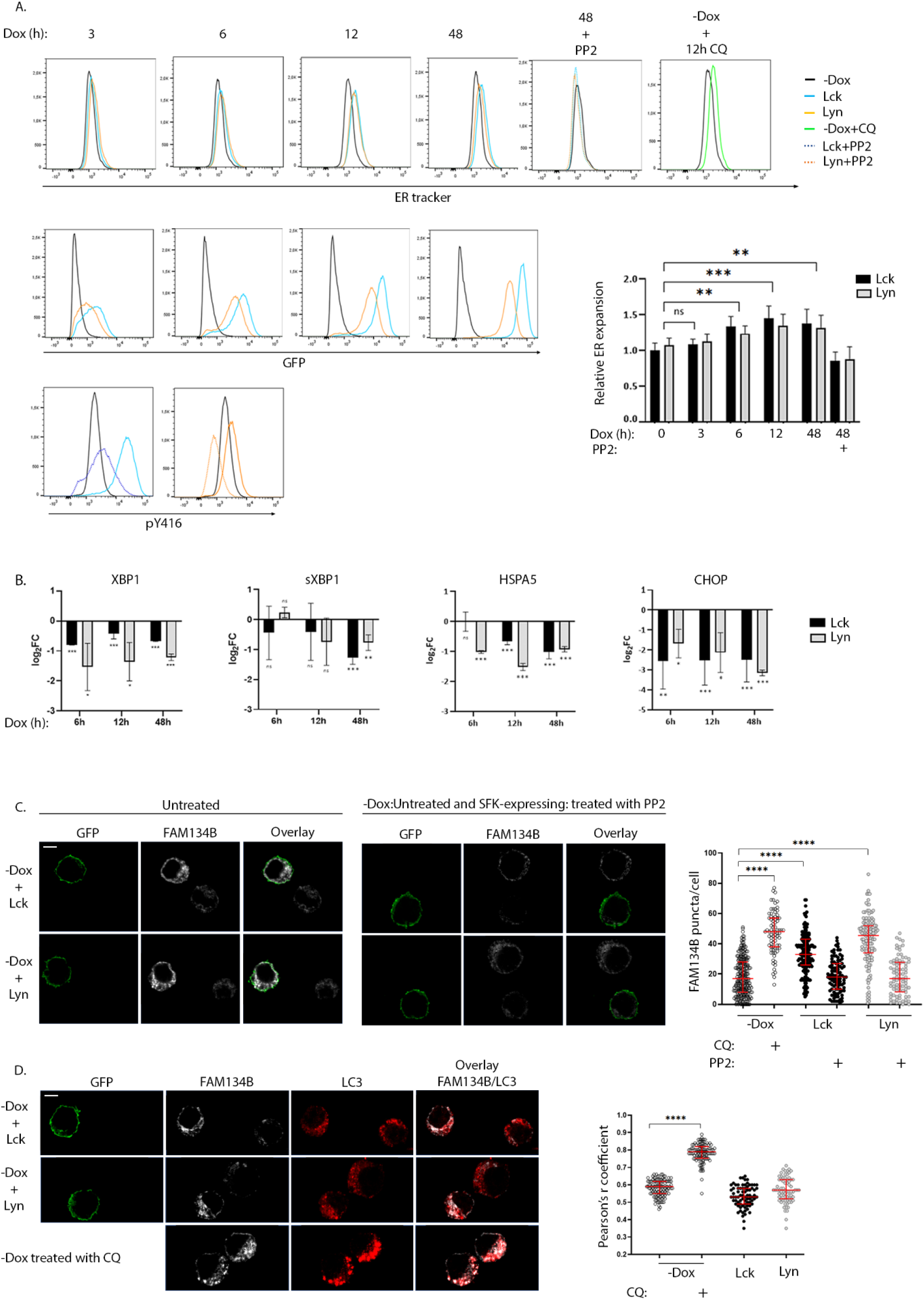
Role of SFKs in the regulation of ER homeostasis. **A. SFK overexpression suffices to induce ER expansion in an enzymatic activity-related fashion**. Lck- or Lyn-expressing cells and the -Dox counterparts were stained with ER tracker Blue-White DPX at the indicated time points after Dox addition to the cell cultures and analysed by FACS. ER expansion measured by increases of ER tracker dye fluorescence (lower panel histograms) is correlated to increases in SFK protein levels (GFP fluorescence, upper panel histograms). 18h incubation with 30μM of the pan SFK inhibitor PP2 reduced ER expansion to baseline (-Dox) levels. The correspondent reduction of pY416 after PP2 incubation for each SFK is shown in the bottom-panel histograms. Treatment with 50 μM of the ER-stress inducer Chloroquine (CQ) for 12h was used a positive control for the expected function of the ER-tracker reagent. Graph shows the ratio of ER tracker dye MFI from Lck- or Lyn-expressing cells to -Dox treated cells. Since essentially there was no difference between the Lck and Lyn samples, statistics were calculated for the mean values between each time point (n=4, Unpaired Student t test; mean +/- SD; **P < 0.01, ***P < 0.001; ns: not significant). **B. SFK-driven ER expansion is not concurrent with upregulation of UPR mediators** RT-qPCR of the indicated UPR modulators. Each time point corresponds to samples stained with ER tracker shown in A. RT-qPCR data are shown log_2_FC of the gene expression between Lck-BJAB (black)/-Dox and Lyn-BJAB (grey)/-Dox (n=2, Multiple unpaired Student t test; mean +/- SD; *P < 0.05, **P < 0.01, ***P < 0.001; ns: not significant). **C. SFK overexpression suffices to induce FAM134B oligomerization in an enzymatic activity-related fashion**. Lck- or Lyn-expressing cells and their -Dox counterparts were mixed at a ratio of 1:1, immobilized on poly-D-lysine coated microscope slides, stained for FAM134B (white), and visualized by confocal microscopy. GFP (green) denotes the SFK-expressing cells. Sample sets include: i) untreated cells (left-hand side panels) ii) SFK-expressing cells treated with 30μM PP2 for 18h, mixed with untreated -Dox cells (right-hand side panels) to provide a comparative visualization of SFK activity influence on the cargo receptor clustering and iii) -Dox cells treated with 50 μM CQ for 12h (indicative image shown in lower panels of D, below). FAM134B oligomers within single cells, appearing as distinctive puncta, were quantified for each sample. Collective quantitation results from three independent experiments (for the untreated samples) and two independent experiments (for PP2 and CQ treatments) are displayed on the adjacent graph. Number of cells for each sample -Dox/untreated n=239, -Dox/CQ n=79, Lck-BJAB/untreated n=164, Lck-BJAB/PP2 n=98, Lyn-BJAB/untreated n=144, Lyn-BJAB/PP2 n=84. Scale bar, 5 μm (n=2, Unpaired Student t test; mean +/- standard deviation [SD]; ****P < 0.0001). **D. SFK-driven FAM134B oligomerization does not coincide with LC3 colocalization**. Lck- or Lyn-expressing cells and their -Dox counterparts mixed at a ratio of 1:1 (upper panels), or -Dox cells treated with 50 μM CQ for 12h (lower panels) were immobilized on poly-D-lysine coated microscope slides, stained for FAM134B (white) and LC3B (red), and visualized by confocal microscopy. GFP (green) denotes the SFK-expressing cells. Colocalization of FAM134B and LC3 was quantified by the coloc2 pre-installed plugin of Image J. Collective colocalization analysis results from two independent experiments are displayed on the adjacent graph. Number of cells for each sample -Dox/untreated n=116, -Dox/CQ n=116, Lck-BJAB/untreated n=76, Lyn-BJAB/untreated n=73. Scale bar, 5 μm (n=2, Unpaired Student t test; mean +/- standard deviation [SD]; ****P < 0.0001).

This discordant profile poses questions as to how ER homeostasis might be maintained in the BJAB-Lck and BJAB-Lyn cell lines. Degradation of excess or damaged ER as well as the turnover of proteasome-resistant proteins by selective autophagy (ER-phagy) is a recently identified mechanism central to ER homeostasis and cellular stress responses (*21*). ER-phagy is dependent on the activation of specialized ER-phagy receptors, few of which were found to be upregulated in our system (Fig. 3D, corresponding heat map). Along these lines, we sought to investigate whether SFK-transduced signals might affect FAM134B, the first identified and so far, the best characterized selective ER-phagy receptor. To execute ER-phagy, FAM134B forms oligomers at cisternae edges of the ER (*22*). This is manifested by the appearance of distinctive FAM134B-positve punctae detected by Immunofluorescence microscopy (*22*). To minimize biases introduced by immunofluorescence staining and/or detection, cells that were cultured in the absence or presence of Dox were mixed at a ratio 1:1 prior to immobilization on microscope slides and staining. Immunofluorescence microscopy using anti-FAM134B antibody revealed enhanced puncta formation of the cargo receptor in the Lck- and Lyn-expressing cells compared to their -Dox counterparts (Fig.4C and Fig.S6A). Importantly FAM134B puncta receded to -Dox levels when the SFK-expressing cells were incubated with PP2 for 18h, (Fig. 4C and Fig. S6A). Progression of ER-phagy is characterized by the recruitment of core autophagy proteins, such as LC3B, to FAM134B clusters and direct association via its LIR motif (*21*). This interaction mediates the recruitment of autophagosomal membranes to the ER and sequestration of ER fragments into LC3B-postive autophagosomes for lysosomal degradation. Accordingly, FAM134B and LC3B have been shown to coprecipitate and colocalize in FAM134B positive puncta (*23*). Unexpectedly, we could not detect colocalization of FAM134B and LC3B in the BJAB-Lck and BJAB-Lyn cell lines (Fig. 4D). Treatment of -Dox cells with Chloroquine induced both FAM134B puncta formation and parallel colocalization with LC3, representative of the canonical ER-phagy pathway (Fig. 4C and 4D).

## Discussion

In this work we examined disruptions in B-cell signaling homeostasis resulting from the overexpression of endogenous (Lyn) and ectopic (Lck) SFKs. Similar to Lck’s behavior in T-cells (*2*), both kinases are constitutively active in B-cells and elicit phosphorylation events in a dose-dependent fashion without any requirement for receptor engagement. However, aberrances in baseline B-cell signaling required the surpassing of a certain expression, and concomitant SFK activity, threshold.

A role for Lck in BCR ITAM phosphorylation has been suggested by studies in CLL cells treated with Lck inhibitors (*24*) and experiments demonstrating co-localization of the kinase with the CD79 chains, under the imposition of BCR ligation (*25*) or in the context of unstimulated subpopulations expressing exceedingly high levels of ectopic Lck (*4*), all of which may represent coincidental events. Here, we demonstrate a direct cause-effect relationship between ectopic Lck and BCR ITAM-driven ignition of signaling responses, which conclude to alterations in the gene expression program. Interestingly, in contrast to Src, Fyn and to a lesser extent Lyn, Lck did not affect the global phosphotyrosine profile of HEK293T cells, indicating a dedication to ITAM/ITIM-containing receptors. This preferential behaviour questions the biological/pathological relevance of ectopic Lck outside the strict leukocyte environment. Both Lck and Lyn overexpression/hyperactivation are considered relevant factors for CLL pathogenesis, and although Lck expression has been linked with resistance to chemotherapy, no association could be found with disease outcome (*26*). Similarly, deletion or expression of hyperactive Lyn in a transgenic mouse model of CLL, despite a clear impact on BCR signaling, failed to establish a connection between the kinase and disease progression (*27, 28*), highlighting the notion that the multifactorial nature of the disease renders difficult the association of pathophysiology with single molecules within signaling pathways.

Comparative analysis between Lck and Lyn designate two parameters governing the threshold of responses downstream antigenic receptors, which are not mutually exclusive. On one hand there is the total amount of SFK activity within the cell, and on the other the individuality of SFK family members, dictated by intrinsic molecular features and/or adaptation to cell-specific regulatory mechanisms. When cells were presented with excessive amounts of active SFKs due to the enforced Lck or Lyn overexpression, unscheduled signaling was initiated and progressed all the way to the nucleus, indicating quantitative overwhelming of the mechanisms designed to keep SFK activity at bay. Nonetheless, for comparable protein levels, Lyn appears more efficient in driving BCR ITAM and downstream effector phosphorylation. This could be due to Lyn displaying higher substrate affinity and/or facilitated juxtaposition to the BCR, dictated by its SH4 and adaptor domains. In parallel however, Lyn is subjected to stricter regulatory restrictions. Lyn exists as two alternatively spliced variants LynA and LynB. LynA is more susceptible to Cbl-mediated degradation; however, our studies show that even the degradation-resistant LynB isoform appears more susceptible than Lck to proteasome-mediated proteolysis and thus not allowed to exceed strictly delimited expression levels. Furthermore, PTP inhibition resulted in the acute augmentation of Lyn’s but not Lck’s activation loop tyrosine phosphorylation. This could be due to a kinetic advantage of Lck trans-autophosphorylation versus dephosphorylation, when compared to Lyn, and/or its preferential segregation in “PTP-free” membrane subregions. The second scenario seems more plausible since Lck pY416 could appreciably increase but required longer pervanadate treatments, implying a delayed rate of encounter between Lck and PTPs. In any case, the time gap of response to pervanadate between Lck and Lyn, implies that the phosphorylation status of Lyn’s activation loop tyrosine is kept under stricter surveillance by PTPs.

Overexpression of both Lck and Lyn generated signaling events capable of altering the gene expression landscape of BJAB cells. We identified a plethora of upregulated mRNAs associated with B-cell activation. Induced genes included HLA molecules, Immunoglobulin constant and variable regions, surface markers and co-receptors, cytokines and their receptors, adhesion and migration molecules, anti-apoptotic mediators and several transcription factors. This induced-gene profile is in accordance with the augmented phosphorylation of proximal BCR signaling transducers and highlights an ability of both SFKs to, at least partially, drive B-cells towards an activation mode in the absence of additional stimulatory signals. Both the GO enrichment analysis and our own comparison of upregulated mRNAs, leaves a general impression that Lck is not as effective as Lyn in the induction of B-cell activation-related genes. This was despite the more robust phorsphorylation of CD79a, Syk, Akt and global phosphotyrosine content recorded in the Lck-BJAB cells. Further studies are required in order to clarify whether this is due to qualitative signaling differences (i.e., incapability of Lck in triggering parallel stimulatory pathways) or due to the more intense activation of SHP-1/CD22 negative regulatory axis and perhaps additional inhibitory pathways. Lyn also triggers both ITAM and ITIM phosphorylation in B-cells, thus simultaneously participates in activatory and inhibitory pathways, but to which side the scales are tipped remains debatable. Lyn-deficient B-cells display hyperresponsive BCR signaling (*29*), implying a net inhibitory effect. On the other hand, expression of a constitutively active Lyn transgene (Y508F) resulted in exaggerated positive proximal BCR signaling (*30*). This phenotype was mimicked by enforced SFK activation via Csk inhibition and revealed that additional B-cell expressed SFKs efficiently triggered inhibitory signaling in the absence of Lyn (*5*). The latter studies suggest a preferential positive influence on B-cell signaling responses. On a similar note, our data showed that the mere elevation of wild-type Lyn expression levels sufficed to elicit proximal BCR signaling and shift the gene expression program towards responses reminiscent of B-cell activation, in the presence of concomitant triggering of the CD22/SHP-1 axis.

A somehow curious observation from the transcriptomics data was the pronounced upregulation of IgHA and IgHE mRNAs and the parallel suppression of AID and accessory DNA repair factors essential for CSR. This was not a peculiarity of the cell line model, since BJAB cells do not constitutively express IgA or IgE isotypes and are used by several groups as a model to study CSR in response to different stimuli. Not surprisingly, many of these reports have documented the concurrence of class switching with AID upregulation (*31*). However, Hauser et al. reported that a-IgM stimulation of mouse splenic B-cells induced to undergo CSR by a 48h LPS/IL-4 treatment, caused sharp downregulation of AID transcripts (*32*). Furthermore, AID gets downregulated as B-cells exit the germinal centers and becomes completely absent upon terminal differentiation (*33*). It was thus proposed that, high-intensity BCR signals shut off AID expression as a means of halting mutagenesis of the antibody genes once good affinity of immunoglobulins has been reached. In our system, cells are in a state of interminable ITAM-transmitted signals, the intensity of which could progressively increase in a time-dependent fashion, in correlation to the ongoing increases in SFK expression levels. The emergence of switched isotype mRNAs inevitably requires AID upregulation at some point, which could possibly have occurred relatively early after SFK expression. Therefore, AID downregulation (detected 48h after Dox addition) may designate a state reminiscent to termination of the switch process. Although based solely on these data, we cannot claim a role of SFKs as tuners of class switching or B-cell terminal differentiation, we strongly believe it is a hypothesis deserving more in-depth exploration.

SFK overexpression affected several genes involved in the apoptotic process. Of note, we noticed parallel upregulation of two members of the BCL2 family with opposing actions, Bcl2 and Bim (BCL2L11). It has been shown in T-cells, that these proteins mutually affect each other’s expression (*34*). Notably, high amounts of Bcl2 increased Bim mRNA levels and was proposed that this codependency has a role in tipping the balance between survival and the timely elimination of activated T-cells in order to restore T-cell homeostasis. Our data implies that a similar mechanism may be in effect for activated B-cells.

Transcriptome analysis of SFK-overexpressing BJAB cells revealed a somehow discordant profile of upregulated Immunoglobulin mRNAs and ER expansion accompanied by marked downregulation of important UPR transducers. To cope with the excessive demand for protein synthesis and folding, antibody-producing B-cells employ mechanisms that increase ER output and alleviate the stress created by the vast accumulation of unfolded/misfolded protein cargo. Accordingly, activation of the UPR signaling pathway drives the upregulation of several target genes that enhance ER functionality, including transcription factors and numerous chaperones and folding enzymes (*35*). An additional mechanism acting in parallel or independently of the typical UPR is the physical expansion of the organelle (*36*). Indeed, activated B-cells can expand their ER volume up to threefold during the differentiation route to plasma cells (*37*). It has been reported that Lyn overexpression can block the induction of CHOP and HSPA5 in airway epithelial cells (*38*). Although in this model Lyn signaling per se did not directly affect the UPR, but rather participated in limiting its intensity under conditions of inflammation-induced ER stress, these data are in line with our observations of a suppressive role of Lyn and Lck in UPR signaling. Lyn-mediated suppression of the UPR in the aforementioned study resulted in inhibition of mucus hypersecretion, it was thus credited with a protective role during allergic reactions in the lung. What might then be the biological value of UPR downregulation in our system? It has been demonstrated that T-cells from an Slfn2 mutant mouse (*elektra* mouse model) display a semi-activated phenotype, which is combined with chronic ER stress and UPR activation, under resting conditions (*39*). As a consequence, *elektra* T-cells, display compromised viability via the intrinsic apoptotic pathway. In this model, knockdown of the major UPR mediators XBP1 or CHOP sufficed to restore viability, thus connecting suppression of the UPR with improved survival. This concept could also apply to our cells. Further to antigen recognition by the BCR, full B-cell activation requires additional signals from the engagement of stimulatory and inhibitory co-receptors, cytokines, and help from T-cells. None of these events take place in our system. Nonetheless, SFK overexpression triggered BCR proximal signaling events and modulated the expression of several genes characteristic of activated B-cells. Based on these facts, one could hypothesize that SFK-overexpressing BJAB cells display what could be characterized as a semi-activated phenotype, which could ultimately compromise their viability as in the case of *elektra* T-cells. In this scenario, the simultaneous UPR downregulation by SFK signaling might serve as a means of preserving survival and reveal a previously unrecognized SFK-mediated mechanism for protecting B-cells from apoptosis under conditions of “inadequate” activating signals. Interestingly, in contrast to Lyn and Lck, the activity of c-Src was essential for UPR activation (XBP1 and HSPA5 induction) in human breast cancer cell lines (*40*) revealing differential involvement of SFKs in stress responses.

An additional noteworthy observation on the transcriptome profiles of SFK-overexpressing BJAB cells was the suppression of genes involved in the Ubiquitin/Proteasome degradation machinery. Assuming that the recorded ER expansion is a consequence of increased protein load delivered to the organelle, one would expect that the proportion of newly synthesized proteins failing to attain their native conformation and thus be destined for degradation should also increase, calling for a reinforcement of the proteasomal machinery. Indeed, augmented protein synthesis is accompanied by a global induction of genes encoding proteasomal subunits (*41*), whereas in yeast, protein misfolding was shown to drive the expression of genes responsible for proteasome biogenesis (*42*). However, a similar to ours paradoxical imbalance between protein production load and cellular degradation capacity has been documented in “short-lived” plasma cells. In this model, differentiating B-cells started downregulating proteasomal activity and subunit expression despite the high rates of Immunoglobulin production. The disproportional proteasomal load to capacity ratio was linked with the onset of apoptosis and was proposed as one of the mechanisms responsible for controlling plasma cell lifespan (*43*). The Ubiquitin-Proteasome system is functionally coupled to the UPR and autophagy (*44*). Inhibition of the former frequently results in compensatory autophagic mechanisms for protein degradation. Our transcriptome analysis revealed induction of several autophagy-related genes implying that SFK-transduced signals favor autophagic catabolic processes in the expense of proteasomal degradation and the UPR. Of particular interest was the induction of genes encoding the ER-phagy receptors RETREG1/FAM134B, CCPG1 and CALCOCO1 and facilitators GABARAP and SEC23. The notion of selective ER-phagy acting as a rheostat between optimal Immunoglobulin production, ER stress responses and viability has been suggested by studies in Atg5 deficient plasma cells (*45*). In this model, autophagic regulation of the ER ensured that Immunoglobulin synthesis, XBP1, HSPA5 expression and ER expansion, would not exceed levels that would be intolerable by the plasma cell.

Our studies revealed oligomerization of the cargo receptor FAM134B under Lck- and Lyn-induced signals. FAM134B oligomerization is mediated via CAMK2B-mediated phosphorylation and signifies activation of the cargo receptor for membrane fragmentation and the onset of ER-phagy (*22*). However, FAM134B activation was not followed by canonical ER-phagy events requiring the recruitment of LC3B for delivery of ER fragments into autolysosomes. Although seemingly counterintuitive, ER-phagy that does not require the core autophagy machinery has been described in yeast (*46*). In this system the process of ER-phagy under stress induced by DTT treatment, was essentially unaffected by the deletion of core autophagy proteins including ATG8 (the yeast ortholog of LC3B). ER-phagy is a relatively newly identified form of autophagy and several aspects of its activation, inhibition and underlying molecular mechanisms remain poorly understood. Nevertheless, emerging evidence support its involvement in numerous human diseases and identify ER-phagy receptors, especially FAM134B, as new therapeutic target (*47*). To our knowledge, this is the first report on the behavior of ER-phagy receptors in B lymphocytes, which prompts for follow-up studies to elucidate the regulation and significance of the very important homeostatic process of ER-phagy in the context of B-cell function under physiological and pathological conditions.

A further point of consideration raised by the transcriptome studies are the numerous DEGs uniquely regulated by Lck and Lyn, which however, could not be assign to unique biological functions by GO analysis. Rather, the two SFKs appear to regulate overlapping pathways albeit with different efficiency. Although further and more detailed studies are required before reaching definitive conclusions, our work implies that SFK-driven tuning of signaling responses may be largely achieved via quantitative rather than qualitative mechanisms dictated by rules of strict enzymatic selectivity.

In conclusion, our model provides an unbiased picture of the signaling machineries set in motion by Lck and Lyn and uncovers several interesting aspects about SFK regulation and function. We show that the WT forms of both kinases are constitutively active in B-cells and can autonomously emit post-translational and transcriptome changes reminiscent of BCR activation. Constitutive ignition of signaling pathways occurred once a certain threshold of SFK expression/activity was surpassed, highlighting limitations in the amplitude of the machinery designed to keep SFKs under control. We further discovered that Lyn was a more effective signal transducer in B-cells and identified specialized control mechanisms designed to keep Lyn, but not Lck, activity levels under strict control. These data may signify that SFKs have been evolutionarily diversified to best suit the needs of the cellular environment they are expressed in and provides an interesting prospect to study whether this trend will be maintained or reversed when Lck and Lyn are overexpressed in T-cells. Finally, we show a previously unrecognized role of Lck and Lyn favoring ER-selective autophagy, which opens new avenues for studying the role of SFKs in the regulation of stress responses especially in the context of lymphocyte differentiation into effector cells.

## Materials and Methods

### Antibodies

Rabbit anti-pSrc (Y416) (2101), anti-pCD79a (Y182) (5173), anti-pSyk (Y525/526) (2711), anti-pSHP1 (Y564) (8849), anti-pAkt (Ser473) (4075), anti-SHP1 (26516), anti-CD22 (98035), anti-β-actin (4970) antibodies were purchased from Cell Signaling Technology; mouse anti-Lck-PE (clone 3A5) (sc-433), anti-CD22 (clone HD39) (sc-73363) antibodies were from Santa Cruz Biotechnology; mouse anti-Phosphotyrosine (clone 4G10) (3560614) antibody was from Merck Millipore; mouse anti-Lyn (610003) antibody was from BD Biosciences; mouse anti-CD79b PE (clone CB3-1) (IM1612U) was from Beckman Coulter. Secondary antibodies for chemiluminescence anti-rabbit and anti-mouse HRP-coupled (7074 and 7076) were from Cell Signaling Technology; secondary antibody for FACS analysis goat anti-rabbit IgG Alexa Fluor 647 (A21245) was from Thermo Fischer Scientific.

### Cloning and plasmids

cDNAs of human wild-type Lck, Lyn isoform B, Fyn and Src were used to generate all plasmid constructs (wild-type, GFP-tagged and mutant LckΔSH4). Vector pcDNA3.1-GFP (Addgene) was used as a template for the subcloning of GFP. Packaging plasmids pVSVG and pSPAX2 were purchased from Addgene. All constructs were cloned in the expression vector pLVX-Tight-Puro (Clontech Laboratories), between 5’ NotI and 3’ EcoRI restriction sites. For the generation of Lck-GFP and Lyn-GFP fusion proteins, GFP was first cloned into pLVX-Tight-Puro between 5’ XbaI and 3’ EcoRI restriction sites. Then, Lck NotI-XbaI and Lyn NotI-XbaI fragments were ligated in-frame, upstream of the GFP sequence in the GFP pLVX-Tight-Puro vector, previously prepared. LckΔSH4 was generated by PCR using a 5’ primer corresponding to amino acids 12-19 of Lck. All constructs were verified by DNA sequencing.

### Cell Cultures and Treatment

Cell lines were maintained at 37°C with 5% CO2 in a humidified incubator (PHCbi). Human embryonic kidney (HEK) 293T (ATCC® CRL-3216) cells were cultured in DMEM (Gibco) supplemented with 10% fetal bovine serum (FBS) (Gibco). JCaM1.6, BJAB (DSMZ ACC-757) and BJAB-transduced cell lines were cultured in RPMI 1640 supplemented with 10% FBS.

For stimulation experiments, cells were incubated with 10 μg/mL of soluble Anti-Human IgM+IgG (eBioscience) at 37°C for 10 min.

For Lck inhibition, cells were incubated with 10 μM of Lck Inhibitor, Lcki (Calbiochem) at 37°C for 10 min. The pan-SFK inhibitor PP2 (Calbiochem) was used at 30 μM for 18h in the humidified incubator (for treatment of samples stained with ER tracker) and at 100 μM for 10 min at 37^°^C (for transient SFK inhibition). For Proteasome inhibition, cells were treated with 1 μM of MG132 (Sigma-Aldrich) at 37°C for different time points. For protein tyrosine phosphatase (PTP) inhibition, cells were treated with 1 mM Pervanadate for 2 or 5 min.

### Pervanadate preparation

Pervanadate solution was freshly prepared before each experiment. Pervanadate stock solution (10 mM) was prepared by adding 100 μl of 100 mM Sodium orthovanadate (Sigma-Aldrich) and 50 μl of 100 mM hydrogen peroxide (diluted from a 3% stock in distilled water) to 850 μl of serum-free RPMI 1640. Pervanadate solution was used within 30 minutes of preparation to minimize decomposition of the vanadate-hydrogen peroxide complex.

### Production of lentiviral particles

Lentiviruses were generated using the packaging cell line HEK 293T. HEK 293T were transfected, at 80% confluency, using Polyethylenimine (PEI, Polysciences) according to the manufacturer’s instructions. The packaging plasmids pVSVG and pSPAX2 were mixed with the lentiviral transfer plasmids containing the gene of interest. PEI solution was added to the plasmid mix, incubated for 20 min at room temperature (RT) and then added to the cells by gently swirling the plate. A complete medium change was performed 18h post transfection. Supernatants containing lentiviral particles were harvested 48h after the medium replacement and immediately used for transduction.

### Transduction and Generation of Tet-On inducible cell lines

Stable, inducible cell lines were generated using the Tet-On Advanced Inducible Expression System (Clontech Laboratories) according to the manufacturer’s instructions. Briefly, 5×10^5^ parental BJAB cells were incubated for 24h with a mix of HEK293T supernatants containing lentiviral particles of the pLVX-Tet-On-Advanced vector (constitutively expressesing the tetracycline-controlled transactivator rtTA-Advanced) and pLVX-Tight-Puro containing the gene of interest, in the presence of 5 μg/ml hexadimethrine bromide (Polybrene, Sigma-Aldrich). After 24h, the transduction supernatants were discarded and replaced with fresh RPMI with 10% FBS and cells incubated for 2 days at 37°C and 5% CO2. Following recovery, the transduced cells were put under selection by Puromycin (5 μg/ml) and Geneticin (200 μg/ml) to generate stable cell line expressing the gene of interest. Expression of the proteins of interest was induced by 3 μg/ml doxycycline (dox, Sigma-Aldrich) added to the culture medium, routinely 48h prior to each experiment.

### Cell Sorting

Before sorting all standard startup, cleaning, and QC procedures, as advised by BD Biosciences, were conducted. BJAB stable cell lines expressing Lck-GFP and Lyn-GFP, were filtered (Falcon 30 μm Cell Strainer, Corning) and 5×10^6^ cells were diluted in 4 ml of FACS buffer (0.5% bovine serum albumin (BSA, AppliChem) in PBS) for immediate sorting. Cells were sorted through the BD FACSAria™ III Cell Sorter (BD Biosciences) using a 70-μm nozzle at event rates up to 10,000 events/s and a system pressure up to 70 psi. The 488-nm and 633-nm lasers were active during sorting procedure on the BD FACSAria™ III Cell Sorter. Standard optical filter and mirror configurations were used for detection of scatter and fluorescence signals. GFP emission was measured using the FITC channel and a cell population with the higher GFP fluorescence was collected (10% of the total GFP positive population). Sorting was conducted at room temperature and sorted cells were collected in 5 ml round-bottom polystyrene tubes (BD Biosciences) containing 500 μl of complete medium (RPMI 1640 supplemented with 10% FBS and 1% penicillin/streptomycin). Immediately after sorting, cells were centrifuged at 1,500 rpm for 5 min, supernatant was discarded, and cells were resuspended and kept in 2 ml of complete medium per million cells overnight (37°C, 5% CO2).

### Flow cytometry

For cell surface staining, cell suspensions were transferred into a 96-well V-bottom plate, centrifuged, resuspended in 100 μl FACS buffer and incubated on ice for 15 min. After centrifugation, supernatants were discarded, and cell pellets resuspended in 30 μl staining solution containing fluorescence conjugated primary antibodies diluted in FACS buffer and incubated on ice for 30 min. Cells were washed twice and analysed on BD Accuri™ C6 Plus Flow Cytometer (BD Biosciences).

For intracellular staining, cells were fixed with BD Phosflow™ Fix Buffer I (BD Biosciences) for 10 min at 37°C. Cells were centrifuged, resuspended in 150 μl permeabilization buffer (0.5% BSA, 0.5% Triton X-100 (CalbioChem) in PBS) and incubated for 15 min at 37°C. Primary antibodies, diluted in wash buffer (0.5% BSA, 0.1% Tween 20 (AppliChem) in PBS), were added to cells and incubated at 37°C for 1h. Cells were then washed twice with wash buffer and incubated with the corresponding secondary antibodies (diluted in wash buffer) at 37°C for 45 min. Samples were washed twice and analysed on BD Accuri™ C6 Plus Flow Cytometer. Acquired data were analysed by FlowJo Software (BD Biosciences). Statistical analysis was performed with Prism (GraphPad Software).

For endoplasmic reticulum staining, live cells were incubated with 250 nM ER-Tracker Blue-White DPX (Thermo Fischer Scientific) for 30 min at 37°C. Cells were then washed once with PBS and analysed on BD FACSCanto™ II Flow Cytometer (BD Biosciences). Acquired data were analysed by FlowJo Software (BD Biosciences). Statistical analysis was performed with Prism (GraphPad Software).

### Immunofluorescence, confocal microscopy image acquisition and analysis

For confocal microscopy sample preparation, single-cell suspension of cells was immobilized on poly-D-lysine (Gibco)-coated coverslips (Marienfeld Superior) for 30 min at 37°C, 5% CO_2_ in a cell culture incubator. Cells were fixed with BD Phosflow™ Fix Buffer I (BD Biosciences) for 10 min at 37°C. Cells were washed once with PBS containing CaCl_2_ and MgCl_2_ (DPBS) and incubated with 1 ml of permeabilization buffer (1% BSA, 0.1% Triton X-100 (CalbioChem) in PBS) for 5 min at 37°C. Cells were washed once with DPBS and incubated with 1 ml of blocking/wash buffer (1% BSA, 0.1% Tween 20 (AppliChem) in PBS) for 10 min at 37°C. Coverslips were then incubated for 1 h at 37°C with the indicated primary antibodies, diluted in wash buffer. Coverslips were rinsed three times and incubated for 1 h at 37°C with corresponding fluorescent secondary antibodies, diluted in wash buffer. After three washes with wash buffer, coverslips were mounted on microscope slides, using one drop of ProLong Gold Antifade Mountant (Thermo Fischer Scientific). Confocal microscopy analyses were carried out in a Leica TCS SP5 confocal scanning microscope (Leica Microsystems) using 488, 561 and 633 nm laser lines and a 63x/1.4 oil immersion lens. Images were acquired in the microscope with a step size of 0.8 μm.

Puncta quantification and colocalization analysis were performed using ImageJ software (National Institutes of Health, available at http://rsbweb.nih.gov/ij/). For better puncta visualization and quantification, acquired images were processed by background subtraction. Puncta were counted manually. Colocalization between LC3B and FAM134B molecules was quantified from the raw images via intensity-based correlation analysis, using the Coloc2 plugin of ImageJ to obtain the Pearson’s correlation coefficient. Region of interest (ROI) around the outer leaflet of the plasma membrane was used on every cell. A single mid-section z-stack was used for each colocalization measurement.

### Immunoprecipitation and Western Blotting

Cells were lysed with ice-cold lysis buffer containing 20 mM Tris-HCl (pH 7.5), 150 mM NaCl, 1% NP-40, 0.5% n-Dodecyl-β-D-maltoside (LM), 1 mM Na_3_VO_4_ and Protease Inhibitor Cocktail 5 (AppliChem) for 15 min on ice. Cell lysates were centrifuged at 14,000g for 15 min at 4°C to remove cell debris. Protein concentration was measured using Bradford solution (AppliChem). Clarified lysates were incubated with the appropriate antibodies for 2h at 4°C and subsequently with protein A/G agarose beads (Santa Cruz Biotechnology) for 1h at 4°C. Immune complexes were washed twice with lysis buffer and boiled in Laemmli buffer at 95°C for 10 min. Then, lysates and immunoprecipitates were separated by SDS-PAGE with a 10% separation gel and transferred to nitrocellulose membranes (GE Healthcare). Following transfer, membranes were blocked for 1h at RT with 5% BSA diluted in TBST (0.05% Tween 20, pH 7.6) and then incubated with primary antibody overnight at 4°C. After washed with TBST, membranes were incubated with HRP–conjugated secondary antibodies for 1h at RT. Detection was performed using Pierce ECL Western Blotting Substrate (Thermo Fischer Scientific) and either developed on Autoradiography Film (Santa Cruz Biotechnology) or scanned using ChemiDoc MP Imaging System (Bio-Rad).

### RNA extraction and Next Generation Sequencing

Total RNA extraction from 2 independent BJAB-Lck, BJAB-Lyn and -Dox cell cultures was performed using TRIzol (Thermo Fischer Scientific) according to the manufacturer’s protocol and was followed by DNase I (NEB) treatment. The quality of the RNA was visualized using agarose gel electrophoresis. Ribosomal RNA was depleted using the RiboMinusTM Eukaryote Kit v2 (Thermo Fisher Scientific). RNAs was quantified using the Qubit RNA HS (High Sensitivity) Assay Kit (Thermo Fisher Scientific) with the Qubit Fluorometer. cDNA libraries were prepared from 100 ng of rRNA-depleted total RNA using the Ion Total RNA-Seq v2 Kit (Thermo Fisher Scientific) according to the manufacturer’s instructions. In brief, RNAs were digested with RNase III and the produced 200nt RNA fragments were successively treated for adapter ligation, reverse transcription, and 14 cycles of PCR amplification using the Ion Xpress™ RNA-Seq Barcode 1-16 Kit (Thermo Fisher Scientific). Yield distribution of the libraries was measured with the Qubit 1x dsDNA HS Assay Kit (Thermo Fisher Scientific) and size distribution was assessed using the Agilent High Sensitivity DNA Kit on the Bioanalyzer 2100 (Agilent Technologies, Santa Clara, CA, USA). Sequencing was performed using an Ion 540™ chip and the 540™ Chef kit on an Ion GeneStudio S5 sequencer.

### Analysis of RNA-seq data

The data were uploaded to the Galaxy server through the usegalaxy.eu public server (*48*). Reads were initially trimmed with trimmomatic (Version 0.38) with the following parameters SLIDINGWINDOW:30:15 MINLEN:25 and mapped on the whole human genome sequence [Genome Reference Consortium Human Build 38; hg38] using the STAR aligner (v2.7.8a) with default parameters except for --sjdbOverhang 75 --outSAMmapqUnique 255 --chimSegmentMin 18 -- chimScoreMin 12 --outFilterType BySJout (*49, 50*). The unmapped reads were subsequently aligned to hg38 using bowtie2 (v2.4.5) using the --very-sensitive-local preset (*51*). The generated alignment files were then merged using picard MergeSamFiles (v2.18.2.1) and the unmapped reads were filtered out using samtools view (v1.15.1) (*52*). Read counts and differential gene expression analysis was performed using featureCounts (v2.0.1) and limma-voom (v3.50.1) respectively (the analysis included genes with counts per million > 0.5 in at least one sample) (*53, 54*). All raw sequencing data are available at the GEO Repository under the Accession Number: GSE212759. The identified differentially expressed genes (DEGs) were subjected for further functional enrichment analysis, using the PANTHER classification platform (*55*). The most representative classes of all the DEGs with log_2_FC <-0.5 and >0.5 are presented in Figure 3C.

### RT-qPCR validation of differentially expressed genes

For quantitative PCR (qPCR), total RNA was extracted from cells by TRIzol reagent (Thermo Fischer Scientific) according to the manufacturer’s protocol and treated with DNase I (NEB). 1 μg of the RNA was reversed transcribed using SuperScript II Reverse Transcriptase (Invitrogen) with random hexamers following the manufacture’s protocol. The expression analysis of the genes was performed using specific primers (Table S1) and normalization was performed based on the expression levels of *ACTB*. qPCR reactions were performed using the KAPA SYBR® FAST qPCR Master Mix (2X) Kit (Kapa Biosystems) on an Agilent Stratagene Mx3000P Multiplex Quantitative PCR System under the following conditions: 95°C for 3 min, followed by 40 cycles of 95°C for 3 s, 60°C for 20 s. The Ct values were extracted and analysed using the 2^−ΔΔCT^ method. The statistical analysis was performed using an independent-samples t-test. All experiments were performed in duplicates.

### Statistical analysis

Statistical analysis was performed using the GraphPad Prism 9 (GraphPad Software). Data were analysed using unpaired two-tailed Student t test or multiple unpaired two-tailed Student t test. All data are presented as mean +/- SD. A p-value less than 0.05 was considered significant (*P < 0.05, **P < 0.01, ***P < 0.001, ****P < 0.0001; ns: not significant). All experiments were repeated with sufficient reproducibility.

## Acknowledgements

We are grateful to Dr D. Efremov (International Centre for Genetic Engineering and Biotechnology, Trieste, Italy) for provision of the parental BJAB cell line.

## Funding

This work was supported by:

Asklepios Program (Gilead Science Hellas) grant (KN)

Fondation Santé grant (KN)

GSRT Research-Create-Innovate grant T2EΔK-00474 (KN)

Andreas Mentzelopoulos Foundation, PhD Scholarship (NK)

## Competing interests

The authors declare that they have no competing interests.

## Data availability

All raw sequencing data are available at the GEO Repository under the Accession Number: GSE212759. All data needed to evaluate the conclusions in the paper are present in the paper and/or the Supplementary Materials.

## Notes

### Competing Interest Statement

The authors have declared no competing interest.

### Summary of Updates

Text in the results section was updated to clarify Figure 4; Figure 4 revised; Supplemental files updated; Methods were moved to main text; Supplementary figure 6 added

