## Supplementary Information for "Integrated signaling and transcriptome analysis reveals Src-family kinase individualities and novel pathways controlled by their constitutive activity"

### Supplemental Text

Development of a B-cell line model for ectopic Lck expression

The successful implementation of our studies required the model B-cell line system to meet the following criteria: i) Absence of endogenous Lck expression, ii) attainment of Lck expression levels adequately high as to trigger autonomous BCR signaling (4) and iii) efficient *ex vivo* activation of BCR signaling pathways by convenient stimulation methods such as a-IgM antibodies. The selection process for developing such a system involved the introduction of Lck in five widely used B-cell lines (MEC-1, IM-9, HS-sultan, Raji and BJAB), and their evaluation to meet the aforementioned criteria. Comparative analysis for Lck and surface BCR expression (Fig. S1A) revealed that, highest expression of Lck was obtained in the Raji and BJAB stable lines. Raji, did not contain appreciable levels of BCR on their cell surface, (Fig. S1B), thus were rendered unsuitable for our studies. BJAB, a Burkit lymphoma cell line that has been widely used to study BCR signaling, met the criteria for high Lck ectopic expression, and robust responses to a-IgM stimulation. Therefore, it was chosen as the best candidate for proceeding with our subsequent analyses.

Incubation with Dox revealed a time-dependent induction of Lck expression with highest levels being achieve after 48h (data not shown), thus all subsequent analyses were performed 48h after Dox addition. Non-Dox treated cultures invariably displayed baseline staining with all antibodies used in this study, and regardless of the constructs they had been transduced with (data not shown). Therefore, for all subsequent analyses, -Dox samples are a mixture of cell lines used for each corresponding experiment.

Fig. S1

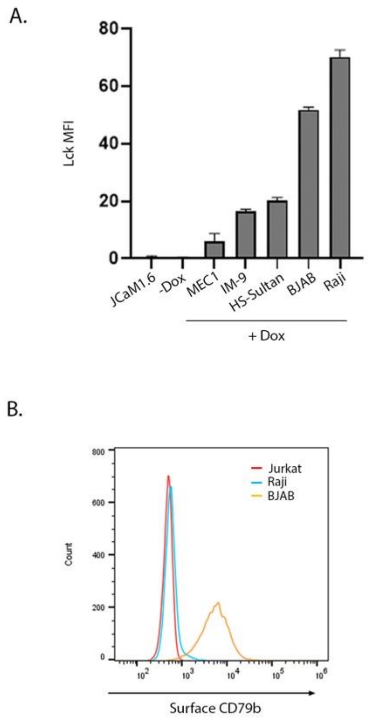


**Fig S1. Development of a B cell model system for ectopic Lck expression**

**A. Comparison of Lck Dox-inducible expression in B cell lines**

The indicated B-cell lines were stably transduced with the lentiviral vector pLVX-Tight-Puro encoding Lck WT. 48h after Dox addition, cells were stained for Lck total protein expression and analysed by FACS. JCaM1.6 cells (an Lck-deficient jurkat variant) were used as a negative control for a-Lck antibody staining. Bar graph indicated a-Lck MFIs (n=2).

**B. The Raji cell line lacks surface BCR expression**

Unfixed live Lck-transduced Raji and BJAB cells were stained CD79b surface expression, Jurkat cells were used as negative control for aCD79b antibody staining.

Fig.S2

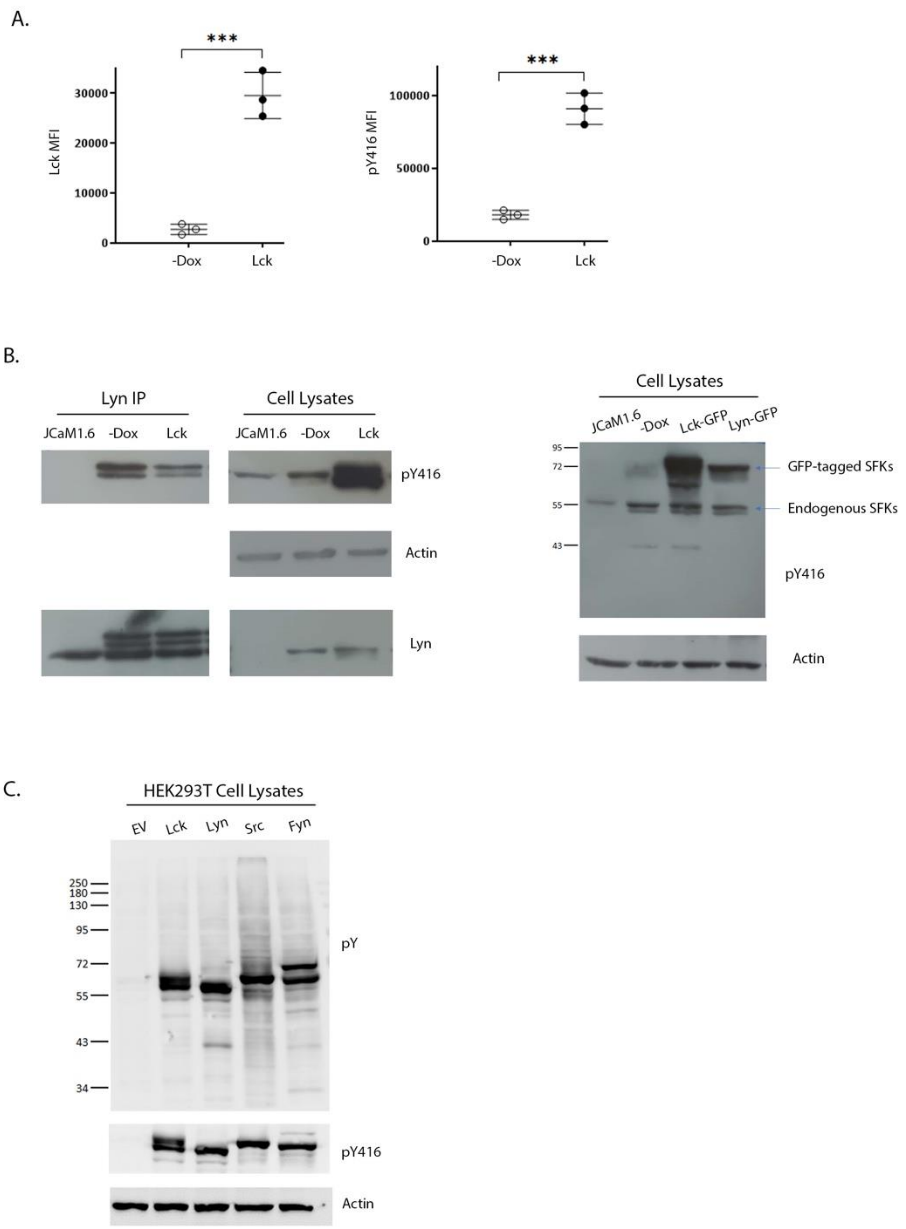


**Fig S2. Profile of cells with ectopic Lck expression**

**A. Ectopic Lck dramatically increases levels of SFK activity in BJAB cells**

Lck-expressing BJAB cells and their non-Dox treated counterparts were stained for Lck and pY416 and analysed by FACS. MFI values for each antibody staining were determined by analysis with the Flowjo software. Graphs show increases in total protein expression and SFK activity levels (n=3, Unpaired Student t test; mean +/- SD; ***P < 0.001).

**B. Ectopic Lck in BJAB does not affect the activation status of Lyn or other B-cell expressed SFKs**

<u>Left-hand side</u>. Endogenous Lyn was immunoprecipitated (IP) from the Lck-BJAB cell line in the presence or absence of Dox. Lyn IPs and total cell lysates were analysed by western blotting with a-pY416 (upper

panels) and a-Lyn (lower panels) antibodies. Actin blot (middle panel) verifies equal sample loading for the cell lysates. JCaM1.6 cells (devoid of both Lyn and Lck expression) were used as a negative control. Increased reactivity of the a-pY416 antibody in the +Dox sample of total Lysates (but not the IPs) is due to ectopic Lck (Fig S2A).

Right-hand side. Total cell lysates from BJAB cells expressing GFP-tagged Lck or Lyn and their non-Dox treated counterparts were analysed by western blotting with a-pY416. The presence of GFP increases the MW of the kinases, thus allowing visualization of both the tagged proteins and endogenous levels of active SFKs (as indicated by the corresponding arrows). Actin blot (lower panel) verifies equal sample loading.

**C. Ectopic Lck does not increase global tyrosine phosphorylation in HEK293T human embryonic kidney cells**

Total cell lysates of HEK293T cells transiently transfected with empty vector (EV) or the indicated SFK members, were analysed by western blotting with the global a-phosphotyrosine antibody, clone 4G10. Loer-panels, a-pY416 and a-actin blots of the same samples verify equal levels of activity amongst individual SFKs and sample loading, respectively.

Fig. S3

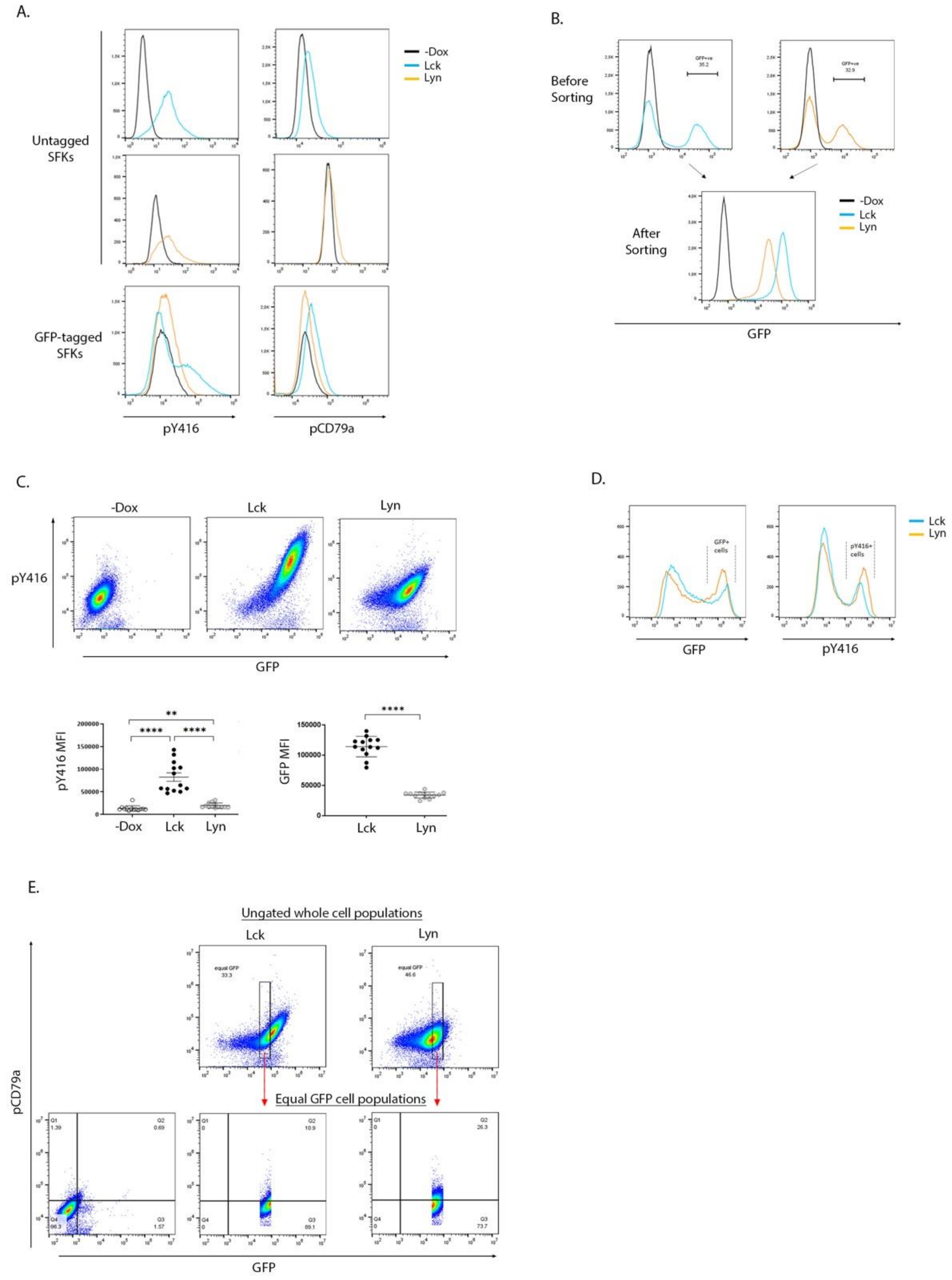


**Figure S3. GFP-tagged SFK behavior and sorting**

**A. Addition of C-terminal GFP tag in Lck and Lyn does not alter their activation and substrate phosphorylation capacity**

Comparative pY416 and pCD79a staining of BJAB cells expressing untagged and GFP-tagged Lck and Lyn as indicated. -Dox samples are used as negative staining controls in each case.

**B. FACS cell sorting of GFP+ populations**

GFP histograms of Lck- and Lyn-expressing cells prior (upper panels) and after (lower panel). Fluorescence activated cell isolation of GFP+ populations (as indicated by the corresponding gates on the upper panels histograms).
Sorted cells were cultured and used for all subsequent experimental analyses.

**C. Lck-BJAB cells have higher levels of GFP expression and consequently SFK activity on a whole cell population basis**

BJAB-Lck and BJAB-Lyn cells were stained with a-pY416 and analysed by FACS in parallel to their -Dox counterparts. 2D FACS plots demonstrate the dose-response relationship between protein expression (GFP) and SFK activity (pY416). Graphs depict MFI values within the whole cell population for each cell line (n=13, Unpaired Student t test; mean +/- SD; **P < 0.01, ****P < 0.0001).

**D. Equivalent levels of Lck and Lyn expression in transfected HEK293T cells**

Histograms of HEK 293T cells were transiently transfected with the indicated constructs and analysed by FACS for GFP and a-pY416 fluorescence. The indicated gates for GFP+ and pY416+ cells were set on empty vector transfected cells.

**E. Equal GFP gating strategy**

Our studies required comparison of Lyn and Lck on an equivalent protein-expression basis. Since it was unattainable to obtain Lyn expression similarly high to Lck in B-cell lines, we resolved in following an "equal GFP gating strategy". In the illustrated example, FACS analysed samples of cells stained for pCD79a, are displayed as 2D FACS plots versus GFP fluorescence. Rectangle gating isolates populations of Lck- and Lyn-expressing cells with equal GFP MFIs (upper panels). These populations are further subdivided by quadrant gates set on the negative control (-Dox) sample (lower panels). The frequency of cells displayed in the upper right quadrant (double-positive for SFK expression and pCD79a) is used for further analysis.

Fig. S4

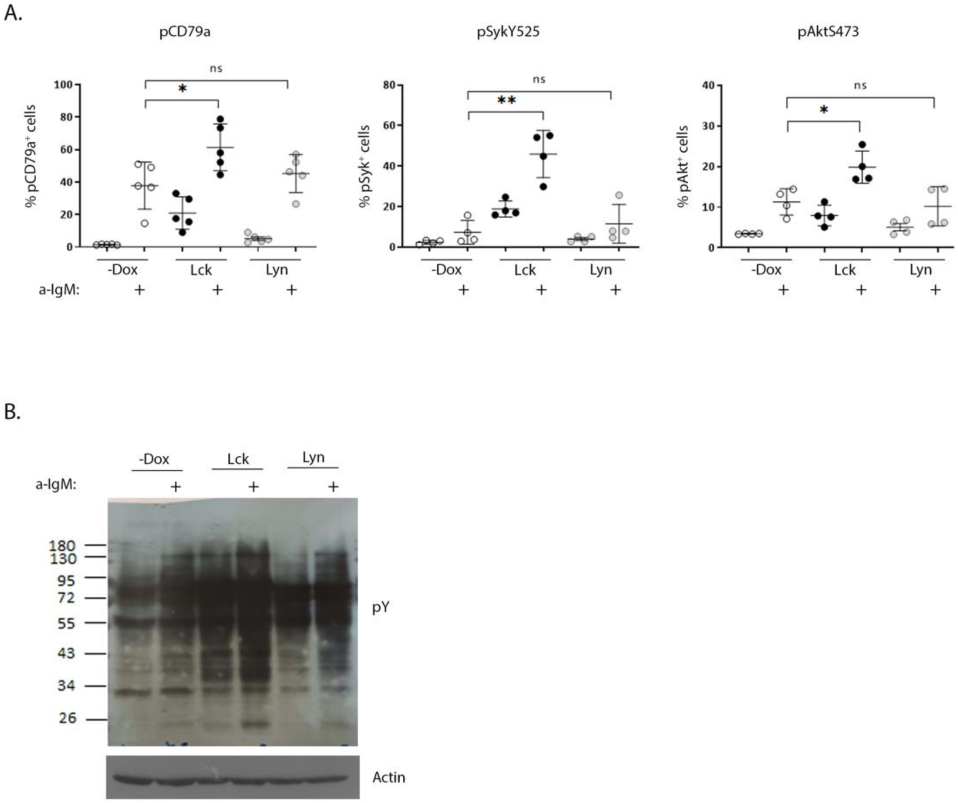


**Figure S4. Lck and Lyn provide a relative advantage in the magnitude of a-IgM triggered responses**
**A**. BJAB-Lck or BJAB-Lyn cells were either left untreated or were stimulated with a-IgM for 10 min at 37ºC prior to staining with the indicated antibodies and analysed by FACS. Graphs depict frequency of cells double-positive for SFK expression and pCD79a for the whole cell population ($n \geq 4$, Unpaired Student t test; mean +/- SD; *$P < 0.05$, **$P < 0.01$; ns: not significant).
**B**. anti-phosphotyrosine (4G10) and actin western blots of total cell lysates of the indicated samples. Stimulation conditions as in A.

Fig. S5

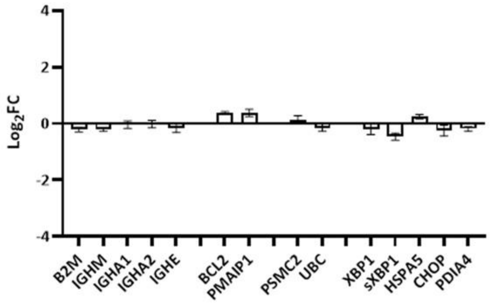


**Fig S5. Doxycycline does not affect representative gene expression in BJAB cells**
RT-qPCR for the validated RNA-seq derived differentially expressed genes shown in Fig. 3E, in parental BJAB cells cultured in the absence or presence of Dox. The y-axis shows the $log_2FC$ of the gene expression of the parental cell line +Dox compared to the same cell line –Dox (n=2, Multiple unpaired Student t test; mean +/- SD, no statistical significance).

Fig. S6

A.

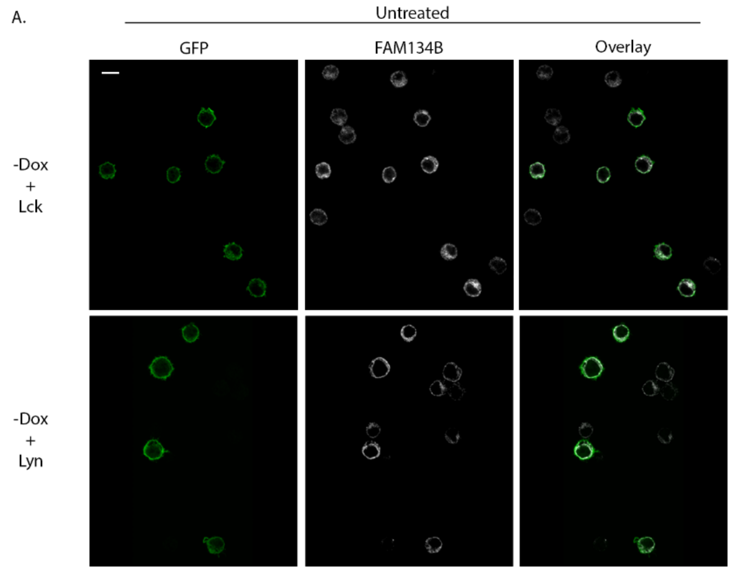
Untreated
GFP
FAM134B
Overlay
-Dox
+
Lck
-Dox
+
Lyn

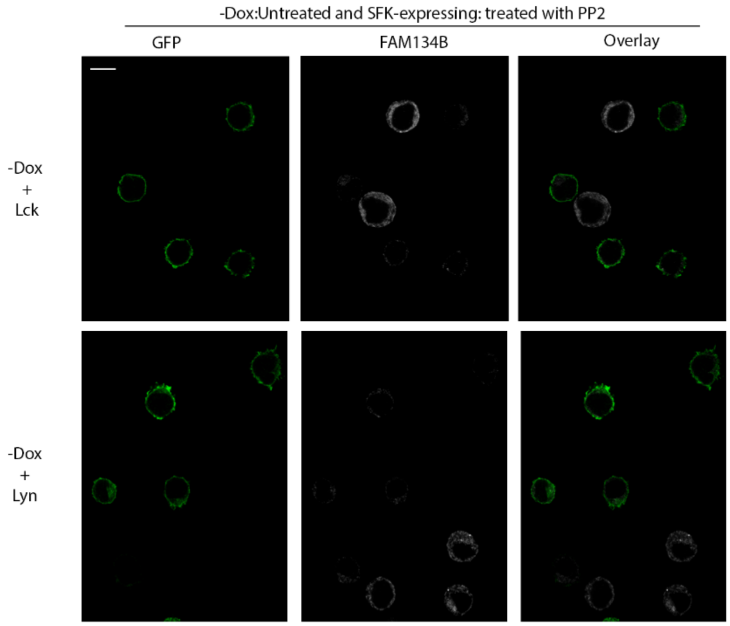
-Dox:Untreated and SFK-expressing: treated with PP2
GFP
FAM134B
Overlay
-Dox
+
Lck
-Dox
+
Lyn

B.

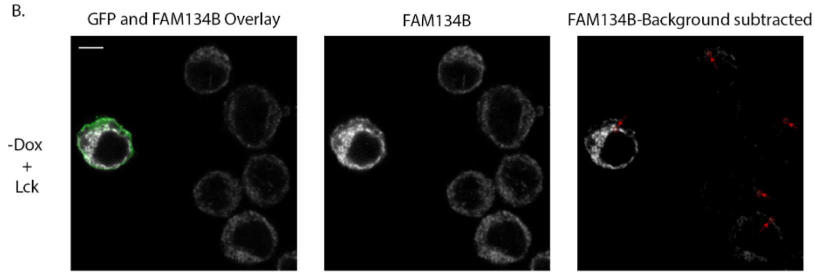
GFP and FAM134B Overlay
FAM134B
FAM134B-Background subtracted
-Dox
+
Lck

**Fig S6. SFK enzymatic activity drives FAM134B oligomerization**

**A.** Low magnification images of samples displayed in Fig. 4C. Untreated cells in upper panels and PP2-treated cells in lower panels as indicated. GFP (green), FAM134B (white). Scale bars, 10 μm.

**B.** To better visualize puncta features and facilitate quantification, acquired images were processed by background subtraction (right-hand side panel). Puncta were counted manually. Examples of what were considered as FAM134B positive puncta in SFK-expressing (GFP positive cells) and in -Dox cells, are indicated by red arrows in the right-hand side panel. Scale bar, 5 μm.

**Table S1**. Specific primers for genes used in the qPCR.

| | Forward primer | Reverse primer |
|---|---|---|
| *B2M* | AGATGAGTATGCCTGCCGTG | TCATCCAATCCAAATGCGGC |
| *HLA-DPB1* | GATGCCTTCAGTCTCCCTGG | GAGCACAGTAGCTTTCGGGA |
| *HSPA5* | CACTCCTGAAGGGGAACGTC | TCAAAGACCGTGTTCTCGGG |
| *DDIT3* | CACCTTTCCCAGAAGTGGCT | TGCGTATGTGGGATTGAGGG |
| *XBP1* | CAGACTACGTGCACCTCTGC | CTGGGTCCAAGTTGTCCAGAAT |
| spliced *XBP1* | GCTGAGTCCGCAGCAGGT | CTGGGTCCAAGTTGTCCAGAAT |
| *PDIA4* | CAACTCAGTTTTGGCGGAGC | TGGCGAACTTCTTCCCACTC |
| *IGHM* | TGACCTTCCAGCAGAATGC | GAAGATGCTGGCAAAGGATG |
| *IGHE* | ACTATGCCACCATCAGCTTG | GTTTTGTTGTCGACCCAGTC |
| *IGHA1* | ATGGGAAGACCTTCACTTGC | TGTTTCCGGATTTTGAGAGG |
| *IGHA2* | AACCATGGGGAGACCTTCAC | CCGGATTTTGTGATGTTGG |
| *UBC* | CCGGGATTTGGGTCGCAG | TCACGAAGATCTGCATTGTCAAG |
| *PSMC2* | TGAGGGGGATATTGCCTTGTT | CAGCCAAATCCCAGAGTGCT |
| *PMAIP1* | CTGCAGGACTGTTCGTGTTC | GCTCGGTTGAGCGTTCTTG |
| *BCL2* | GGGATTCCTGCGGATTGACA | AATGAATCAGGAGTCGCGGG |
